# Microtubule deacetylation drives kinesin-1 mediated mitochondrial transport accelerating breast cancer cell migration

**DOI:** 10.1101/2025.10.29.685374

**Authors:** Morgane Morin, Sylvie Rodrigues-Ferreira, Cynthia Seiler, Emna Ouni, David Mazaud, Hadia Moindjie, Christian Poüs, Charlotte Aumeier, Kristine Schauer, Clara Nahmias

**Affiliations:** Université Paris-Saclay, Gustave Roussy, Inserm U981, Prédicteurs moléculaires et nouvelles cibles en oncologie, F-94805, Villejuif, France; Inovarion, 75005, Paris, France; Université Paris-Saclay, Gustave Roussy, Inserm U1279, Dynamique des cellules tumorales, F-94805, Villejuif, France; Plateforme Imagerie PICT-IBiSA, Institut Curie, PSL Research University, Paris 75005, France; Institut Curie, PSL Research University, Sorbonne Université, CNRS, UMR3664, 26 rue d’Ulm, Paris 75005, France; INSERM UMR-S 1193, UFR de Pharmacie, University Paris-Saclay, Orsay, France; Laboratoire de Biochimie-Hormonologie, Hôpital Antoine Béclère, AP-HP, Hôpitaux Universitaires Paris-Saclay, Clamart, France; Department of Biochemistry, University of Geneva, 1211, Geneva, Switzerland

## Abstract

Mitochondrial trafficking is reprogrammed in metastatic breast cancer cells to sustain their migratory and invasive behavior. Mitochondria repositioning to sites of high energy demand is governed by a balance between opposing dynein and kinesin-1 (KIF5B) molecular motors whose regulation remains incompletely understood. Here, we identify the *SYBU* gene as a candidate prognostic marker downregulated in metastatic disease. *SYBU* encodes syntabulin, a mitochondria outer membrane protein that interacts with dynein to counterbalance KIF5B-dependent anterograde transport to the cell cortex. Loss of *SYBU* disrupts the balance, causing excessive KIF5B-driven mitochondria movement, microtubule damage and deacetylation. In turn, microtubule deacetylation reinforces KIF5B-mediated transport, creating a positive feedback loop that drives mitochondria distribution close to the cell periphery and enhances cancer cell migration. Pharmacological inhibition of the tubulin deacetylase HDAC6 restores mitochondrial positioning and reduces cell migration in *SYBU*-deficient cells. Our findings identify *SYBU* as a key regulator of mitochondrial trafficking and pave the way to personalized therapeutic approaches for metastatic breast tumors with low *SYBU* expression.

## Introduction

Breast cancer remains a leading cause of death by malignancy among women worldwide. Distant metastasis, which develops in nearly one third of patients, is a fatal complication of the disease (Sung et al., 2021). In the era of personalized medicine, new prognostic biomarkers that stratify breast cancer patients at risk to metastasize are urgently needed. Metastasis is a multistep process in which cancer cells escape from the primary tumor, invade surrounding tissues to reach the blood vessels and ultimately colonize a secondary organ. Central to the metastatic cascade is cancer cell migration, a highly energy-demanding process that heavily relies on mitochondrial function. Proper positioning of mitochondria at the leading edge to provide localized energy supply has emerged as a critical requirement to sustain directional migration (Desai et al., 2013; Caino et al., 2015; Cunniff et al., 2016; Schuler et al., 2017; Daniel et al., 2021). Notably, mitochondrial proteins and molecular motors that orchestrate mitochondrial intracellular transport along microtubules have been tightly associated with cell migration and cancer metastasis (Caino et al., 2016; Altieri, 2019; Furnish and Caino, 2019; Morin et al., 2022).

Mitochondrial transport relies on bidirectional movement along microtubules. Anterograde transport toward microtubule plus ends at the cell periphery is primarily mediated by kinesin-1 motors (KIF5B), which couple to mitochondria *via* TRAK adaptors and the outer membrane GTPases Miro1 and Miro2 (MacAskill and Kittler, 2010; Tang, 2016; López-Doménech et al., 2018). Retrograde transport toward the microtubule minus ends near the nucleus is driven by the cytoplasmic dynein-dynactin complex (López-Doménech et al., 2018). A balance of forces is generated by these opposing motors (Hendricks et al., 2010; Hancock, 2014; Belyy et al., 2016; Rezaul et al., 2016). Disrupting the equilibrium between dynein and kinesin forces in cancer cells likely drives mitochondria mispositioning, yet the key regulators of this balance remain poorly characterized.

Microtubule post-translational modifications (PTMs) provide an additional regulatory layer in intracellular transport. Acetylation of α-tubulin at lysine 40 inside the microtubule lumen has been associated with kinesin-1 binding, motility and anterograde transport (Reed et al., 2006; Ravindran et al., 2017; Tas et al., 2017; Monteiro et al., 2023), although this remains a matter of debate (Walter et al., 2012; Kaul et al., 2014; Balabanian et al., 2017; Verhey and Ohi, 2023). The pattern of microtubule acetylation results from a balance between the activities of tubulin acetyltransferase αTAT1 and tubulin deacetylase HDAC6. Both enzymes enter the microtubule lumen *via* sites of microtubule damage or by their dynamic ends (Janke and Montagnac, 2017). Kinesin-1 activity itself was shown to cause friction forces that create microtubule damage (Triclin et al., 2021; Budaitis et al., 2022). This generates entry sites for HDAC6 inside the lumen, ultimately resulting in a gradient of microtubule deacetylation from the perinuclear region to the cell cortex (Andreu-Carbó et al., 2024, 2025). The interplay between microtubule stability and molecular motor activity has emerged as a key regulator of intracellular organelle transport. Whether mitochondrial proteins themselves finely tune the dialogue between microtubule acetylation and motor movement to govern mitochondrial transport remains an open question.

Syntabulin, the major product of the *SYBU*/GOLSYN gene, is a mitochondrial outer membrane protein originally identified in neurons (Su et al., 2004). This 663-amino acids protein connects mitochondria to microtubules and functions as a kinesin-1 (KIF5B) adaptor in axons, facilitating the anterograde transport of syntaxin (Su et al., 2004) and mitochondria (Cai et al., 2005) along microtubule tracks toward the synapse. To date, the role of syntabulin in cancer remains unexplored.

We show here that *SYBU* expression is markedly downregulated in high-grade, triple-negative breast cancers and in metastatic tissues, and that low *SYBU* levels are associated with poor clinical outcome for the patients. Mechanistically, we show that syntabulin interacts with dynein and attenuates Miro1/KIF5B-mediated anterograde transport. Loss of *SYBU* increases KIF5B-driven mitochondria movement and causes microtubule deacetylation. This in turn amplifies KIF5B-dependent anterograde transport, establishing a positive feedback loop that promotes mitochondrial positioning at the leading edge. We propose that downregulation of *SYBU* triggers accelerated cancer cell migration by orchestrating a crosstalk between microtubule acetylation and KIF5B-driven mitochondria transport.

## Results

### *SYBU* expression is downregulated in breast cancer patients

To investigate the clinical relevance of *SYBU* in breast cancer, we analyzed its expression in primary tumors, metastatic lesions, and normal breast tissues using the publicly available TNMplot database. We observed a marked reduction in *SYBU* expression in breast tumors: 55.3% of primary tumors displayed lower *SYBU* levels compared to normal tissues, and expression was further diminished in 59.8% of metastatic samples (Fig. 1A; Supplementary Table S1). Importantly, this downregulation was not restricted to breast cancer; *SYBU* was also consistently decreased across multiple cancer types, including adrenal, bladder, liver, lung, pancreas, skin, stomach, testis, thyroid, and uterus (Supplementary Fig. S1A).

**Figure 1.**
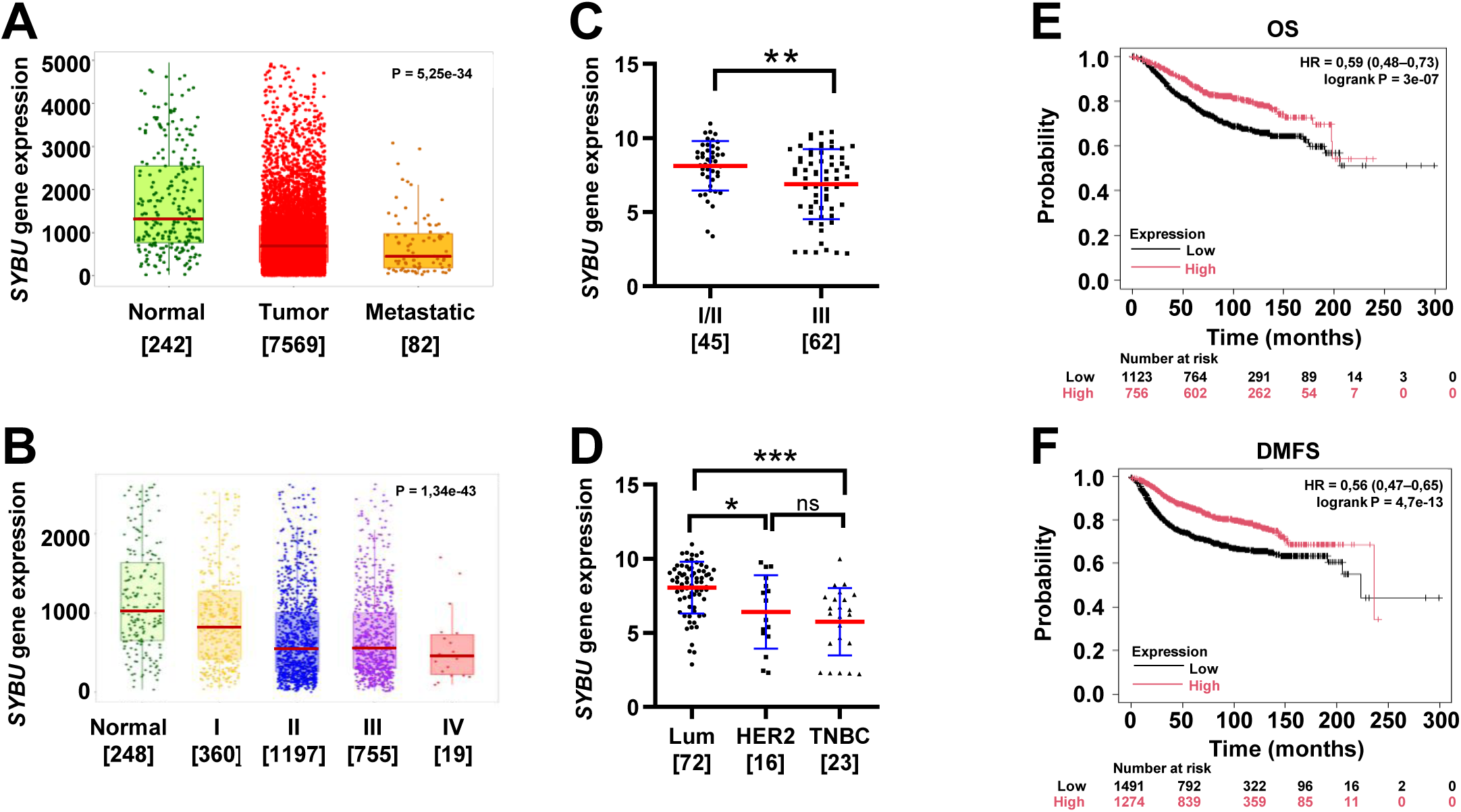
*SYBU* is downregulated in aggressive breast cancer and associated with poor clinical outcome. **(A)** Microarray analysis of *SYBU* expression in normal, tumoral and metastatic breast tissues. **(B)** *SYBU* expression across clinical stages of breast cancer compared to normal tissues. **(C,D)** *SYBU* expression (probeset 218692_at) according to tumor grade **(C)** and molecular subtype **(D)** in the R02 cohort. **(E,F)** Kaplan-Meier survival curves based on *SYBU* expression (probeset 218692_at) showing overall survival (OS) **(E)** and distant metastasis-free survival (DMFS) **(F)**. SYBU high and low expression groups were defined based on the probeset best cutoff. **<u>Data information.</u>** The number of tumors is indicated under brackets. **(A,B)** data are from tnmplot.com; **(E,F)** from kmplot.com. Statistical analyses: **(A,B,D)** Kruskal-Wallis with Dunn’s *post hoc* test; **(C)** Mann-Whitney test; **(E,F)** log-rank test. p*<0.05, p**<0.01, p***<0.001.

Transcriptomic profiling of independent breast cancer cohorts revealed that *SYBU* downregulation correlates with aggressive disease features. Tumors of advanced stage (Fig. 1B) and high grade (Fig. 1C; Supplementary Fig. S1B-D) exhibited significantly lower expression levels compared with their low-stage and low-grade counterparts. Subtype-specific analysis further showed that HER2-positive and triple-negative breast cancers (TNBC), as well as estrogen receptor (ER)-negative tumors expressed significantly less *SYBU* than luminal ER-positive tumors (Fig. 1D; Supplementary Fig. S1E-G).

Interestingly, genetic analyses revealed that *SYBU* is rarely mutated, with somatic alterations occurring in only 0.4% of breast cancer samples (Supplementary Fig. S1H). This strongly suggests that *SYBU* dysregulation in breast cancer occurs predominantly at the transcriptomic level. Finally, survival analyses underscored the clinical significance of *SYBU* expression. Kaplan-Meier curves demonstrated that low *SYBU* expression is significantly associated with worse patient outcomes, including poor overall survival and distant metastasis-free survival (Fig. 1E, 1F), as well as reduced relapse-free survival (Supplementary Fig. S1I).

In summary, these findings establish *SYBU* as a consistently downregulated gene in breast cancer, particularly in aggressive and metastatic tumors. Low *SYBU* expression correlates with unfavorable prognosis, positioning *SYBU* as a potential prognostic biomarker and a candidate for further investigation in breast cancer progression.

### *SYBU* depletion increases breast cancer cell migration

We investigated whether the *SYBU*-encoded protein syntabulin may play a role in regulating breast cancer cell migration, which is an essential step in the metastatic process. We analyzed *SYBU* expression in a panel of breast cancer cell lines (Supplementary Fig. S2A) and we selected triple-negative metastatic D3H2LN breast cancer cells for functional studies, based on their well-established pro-migratory behavior (Molina et al., 2013; Rodrigues-Ferreira et al., 2020). *SYBU* depletion in D3H2LN (Supplementary Fig. S2B, S2C) increased cell migration as assessed by wound healing assay (Fig. 2A). Similar results were obtained in CAL120 and MDA-MB-231 metastatic breast cancer cells (Supplementary Fig. S2B, S2D-S2F). Conversely, expression of GFP-tagged syntabulin (GFP-SYBU) significantly reduced wound closure in control cells and successfully rescued the enhanced migratory phenotype observed in *SYBU*-deficient D3H2LN cells (Fig. 2B). *SYBU* depletion also led to increased chemotactic cell migration in a Boyden chamber assay (Fig. 2C) as well as increased invasion of D3H2LN cells cultured as 3-dimensional multicellular spheroids in a matrigel matrix (Fig. 2D), with no significant effect on spheroids growth (Supplementary Fig. S2G).

**Figure 2.**
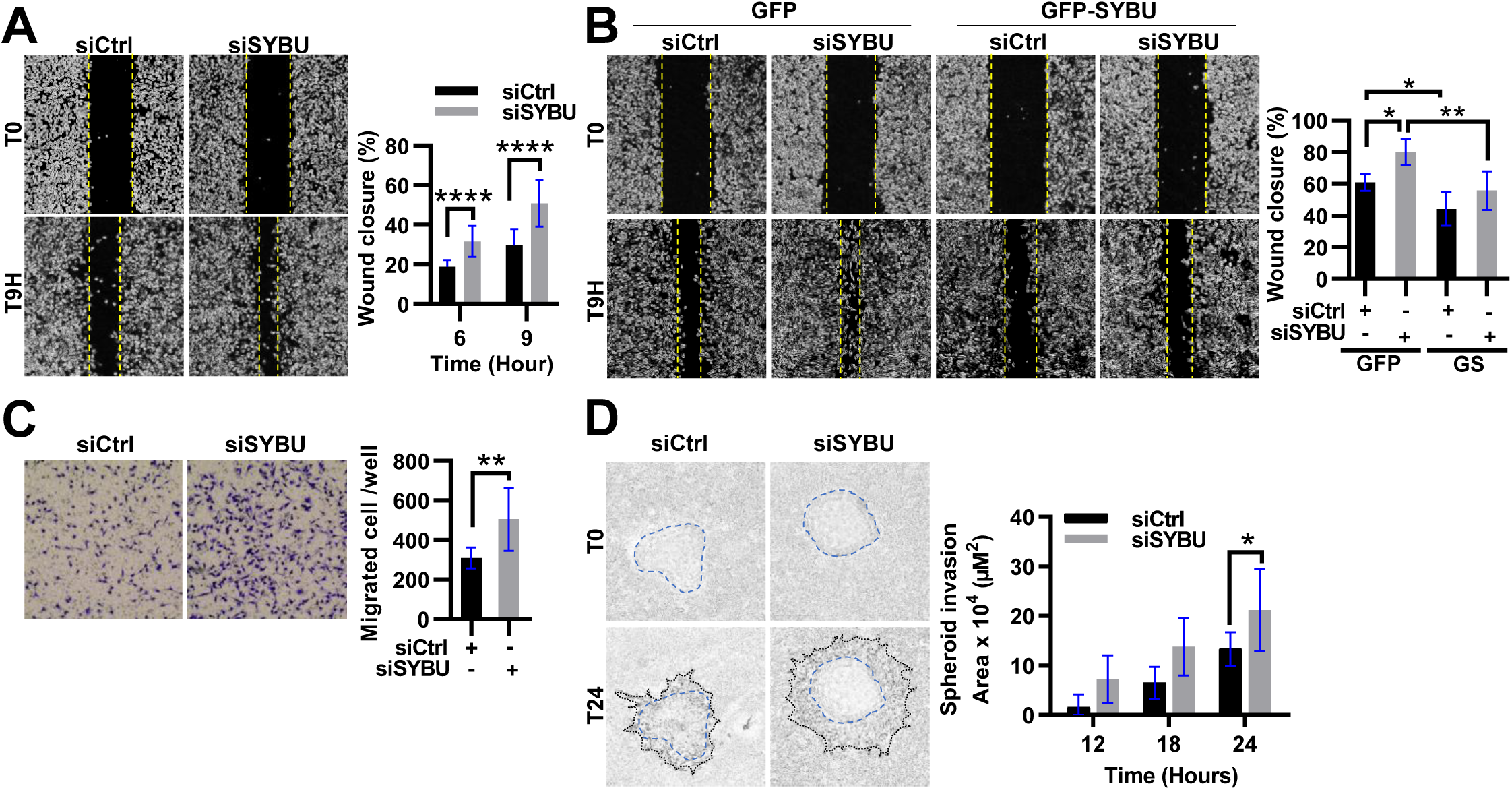
SYBU silencing impairs tumor cell migration and invasion. **(A)** Wound healing assays in D3H2LN cells transfected with control siRNA (siCtrl) or siRNA targeting syntabulin (siSYBU). *Right:* Quantification of wound closure over time. **(B)** Same as (A) in cells transfected with GFP-SYBU or GFP as a control. *Right:* Quantification of wound closure at T9H. **(C)** Chemotaxis assay in siCtrl or siSYBU D3H2LN cells using Boyden chambers. *Right:* Quantification of the number of cells migrating through the membrane after 6 hrs. **(D)** Spheroid invasion assay in siCtrl or siSYBU-transfected D3H2LN cells included in Matrigel (IncuCyte). *Right:* Quantification of area over time. **<u>Data information</u>**. Error bars represent mean ± SD from four **(A)** and three **(B, C)** independent experiments. n=5 wells **(A,B)** and n=3 wells **(C)** per experiment. **(D)** shown is one out of two experiments (n=6 wells). Statistical analyses: **(A)** One-way ANOVA with Tukey’s *post hoc*; **(B)** Kruskal-Wallis with Dunn’s *post hoc*; **(C)** Unpaired T-test and **(D)** Two-way ANOVA with Sidak’s multiple comparisons tests. p*<0.05, p**<0.01, p***<0.001, p****<0.0001.

Together, these results indicate that low *SYBU* expression in breast cancer cells is associated with increased invasive and pro-migratory properties, which is consistent with increased metastatic ability.

### *SYBU* depletion drives anterograde mitochondrial transport

Cell migration is a high energy-demanding process that requires the positioning of mitochondria at the leading edge to supply focal adhesions with the ATP needed for motility (Caino et al., 2015; Cunniff et al., 2016; Daniel et al., 2021). Syntabulin regulates anterograde mitochondrial transport along microtubules in neurons (Cai et al., 2005; Morin et al., 2022). This led us to investigate whether it may also contribute to mitochondrial transport in migrating cancer cells. In *SYBU*-depleted D3H2LN cells undergoing migration, mitochondria localized within membrane protrusions and in close proximity to the tips of microtubules, whereas in control cells they remained predominantly perinuclear (Fig. 3A). In non-migrating HeLa cancer cells, *SYBU* depletion – using two different siRNAs - favored mitochondria positioning close to the cell cortex, visualized by actin staining (Supplementary Fig. S3A-S3E). Super-resolution STORM microscopy further confirmed that in resting *SYBU*-depleted cells, mitochondria are consistently positioned near the cell periphery contrary to *SYBU*-expressing cells (Fig. 3B). *SYBU* silencing also induced peripheral distribution of mitochondria in breast cancer D3H2LN and CAL120 cells (Supplementary Fig. S3F, S3G). Expression of GFP-SYBU was sufficient to rescue the phenotype and restore perinuclear mitochondria distribution in *SYBU*-silenced cells (Fig. 3C). Finally, to quantitatively assess mitochondrial localization in cells of the same shape and size, we used crossbow-shaped micropatterns. Results showed that mitochondrial signal intensity, measured within a well-defined region located at the crossbow tip distant from the nucleus, was elevated following *SYBU* depletion (Fig. 3D), confirming increased peripheral distribution of mitochondria. Furthermore, in *SYBU*-depleted cells, mitochondria were distributed in closer proximity to focal adhesions at the cell periphery (Fig. 3E) suggesting that they may be more efficient in providing energy to these structures that are essential for cell migration.

**Figure 3.**
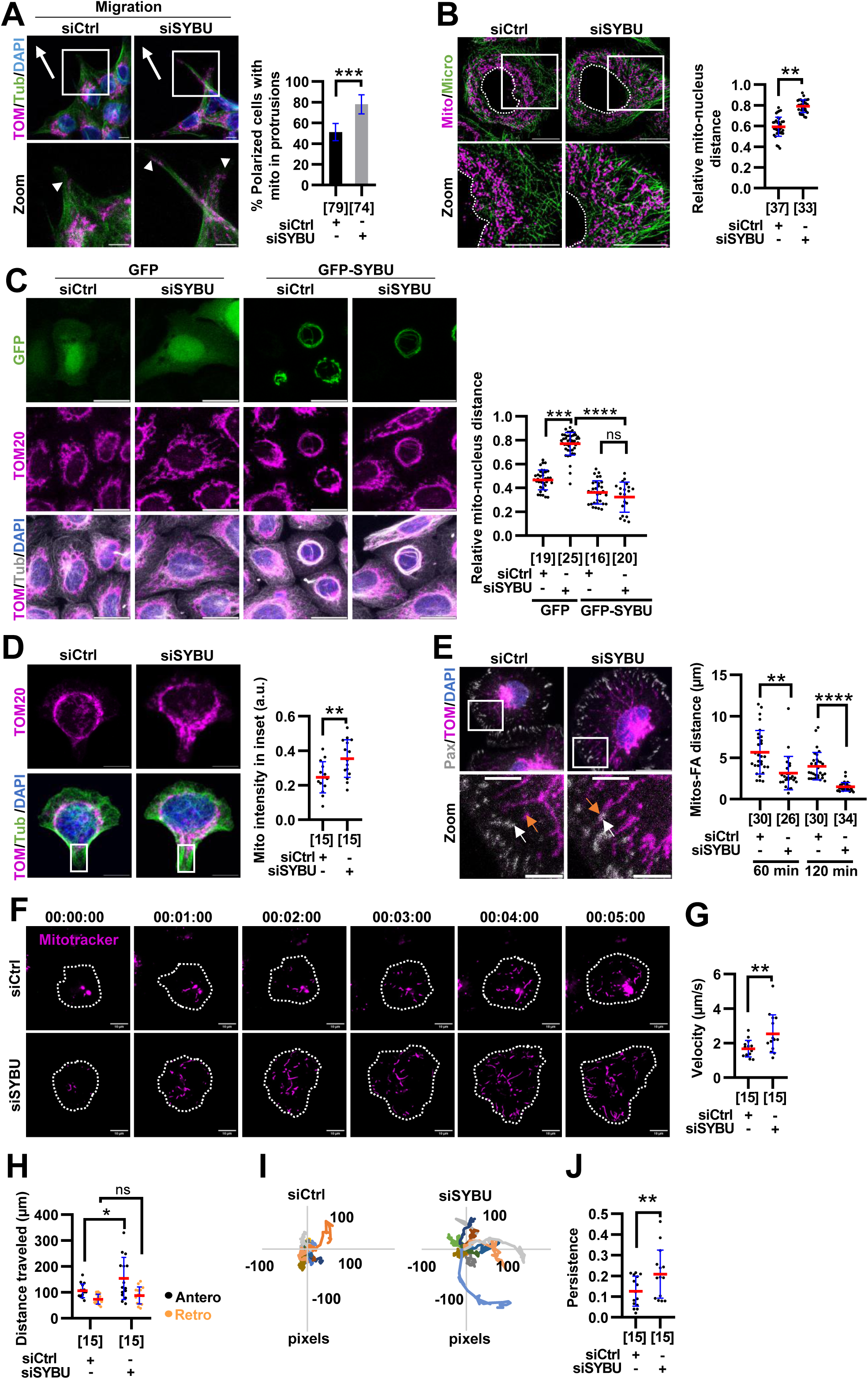
*SYBU* depletion enhances anterograde mitochondrial transport along microtubules. **(A)** Immunofluorescence of mitochondria distribution in siCtrl or siSYBU-transfected D3H2LN cells allowed to migrate for 9 hrs. Arrows indicate the direction of migration. Arrowheads indicate the presence of mitochondria in polarized cells. *Right:* Percentage of cells with mitochondria infiltrating protrusions, normalised to number of polarized cells. **(B)** Super resolution STORM microscopy of mitochondria distribution in resting, siCtrl or siSYBU-transfected, HeLa cells. *Right:* Quantification of relative distance between mitochondria and nucleus. **(C)** Immunofluorescence of mitochondria distribution in siCtrl or siSYBU-transfected HeLa cells expressing GFP-SYBU or GFP as a control. *Right:* Quantification as (B). **(D)** Immunofluorescence of mitochondria distribution in resting siCtrl or siSYBU-transfected HeLa cells grown on crossbow micropatterns. *Right:* Quantification of mitochondria staining intensity in the inset. **(E)** Immunofluorescence of siCtrl or siSYBU-transfected D3H2LN cells during cell adhesion (120 minutes post-seeding). The white arrow shows the position of a focal adhesion (FA). The orange arrow shows the position of the nearest mitochondria. *Right:* Quantification of the distance between FA and the nearest mitochondria (60 and 120 minutes post-seeding). **(F)** Time-lapse imaging showing mitochondria movements (Mitotracker) in siCtrl or siSYBU-transfected D3H2LN cells during cell adhesion (30 minutes post-seeding). Time is indicated in minutes. **(G)** Quantification of mitochondria velocity. **(H)** Quantification of cumulated distance in anterograde (antero) and retrograde (retro) directions. **(I)** Tracking of mitochondria trajectories. **(J)** Quantification of mitochondria persistence. **<u>Data information.</u>** Fluorescent staining shows mitochondria in magenta. Microtubules in green **(A,B,D)** or in grey **(C)**. Focal adhesions (FA) are stained with paxillin in grey **(E)**. Error bars represent mean ± SD from three **(B,D,E)** and two **(C)** independent experiments. **(A)** One representative out of 3 independent experiments. Statistical analyses: **(A,J)** Mann-Whitney; **(B,D,G)** Unpaired T-test; **(C,E)** Kruskal-Wallis with Dunn’s *post hoc* and **(H)** Two-way ANOVA with Sidak’s *post hoc* tests. p*<0.05, p**<0.01, p***<0.001, p****<0.0001. Number of cells is indicated under brackets in **(A-E)**. For **(G-J)**, number of mitochondria is under brackets, n=15 mitochondria tracked in 3 individual cells. Scale bar = 10 µm; zoom = 1 µm **(A)** and 5 µm **(E)**.

Finally, time-lapse videomicroscopy was undertaken to evaluate the parameters of mitochondria transport. Results revealed that mitochondria move 1.5-fold faster in *SYBU*-silenced cells compared to control (Fig. 3F, 3G, Supplementary video 1, video 2). They also cover longer distances in the anterograde direction, while distances traveled in the retrograde direction remain largely unchanged (Fig. 3H). Furthermore, mitochondria displayed a more persistent trajectory and improved directionality in *SYBU*-depleted cells (Fig. 3I, 3J), accounting for peripheral distribution of mitochondria.

Taken together, these findings indicate that in cancer cells, low *SYBU* expression drives mitochondria anterograde transport toward the cell periphery.

### Syntabulin interacts with dynein light chain 1 DYNLL1

We sought to investigate the molecular mechanism by which syntabulin controls mitochondrial transport in cancer cells. The observation that in cells expressing GFP-SYBU, mitochondria are positioned in a perinuclear area raised the possibility that syntabulin uses dynein, the molecular motor responsible for retrograde mitochondrial transport.

Syntabulin is a 663-amino acids protein anchored in the mitochondrial outer membrane through two C-terminal transmembrane α helices (Supplementary Fig. S4A). It contains a coiled-coil domain that interacts with microtubules and a large intrinsically disordered region in its N-terminal portion.

We focused on dynein light chain DYNLL1 (LC8) as a candidate partner of syntabulin as this protein is a hub that contributes to dynein motor function and binds to a panel of proteins with intrinsically disordered regions (Chen et al., 2009; Jespersen et al., 2019; Reardon et al., 2020). Co-immunoprecipitation experiments conducted in cells expressing GFP-SYBU revealed the presence of DYNLL1 in molecular complexes containing GFP-SYBU but not GFP alone (Fig. 4A). Notably, in these experiments no interaction could be detected between GFP-SYBU and the kinesin-1 motor protein KIF5B (Fig. 4A). Proximity ligation assay (PLA) further confirmed spatial proximity between GFP-SYBU and DYNLL1 (Fig. 4B, Supplementary Fig. S4B-S4D). Of note, PLA rolling circles were localized along microtubules (Fig. 4B), indicating that syntabulin and dynein predominantly interact along microtubule tracks.

**Figure 4.**
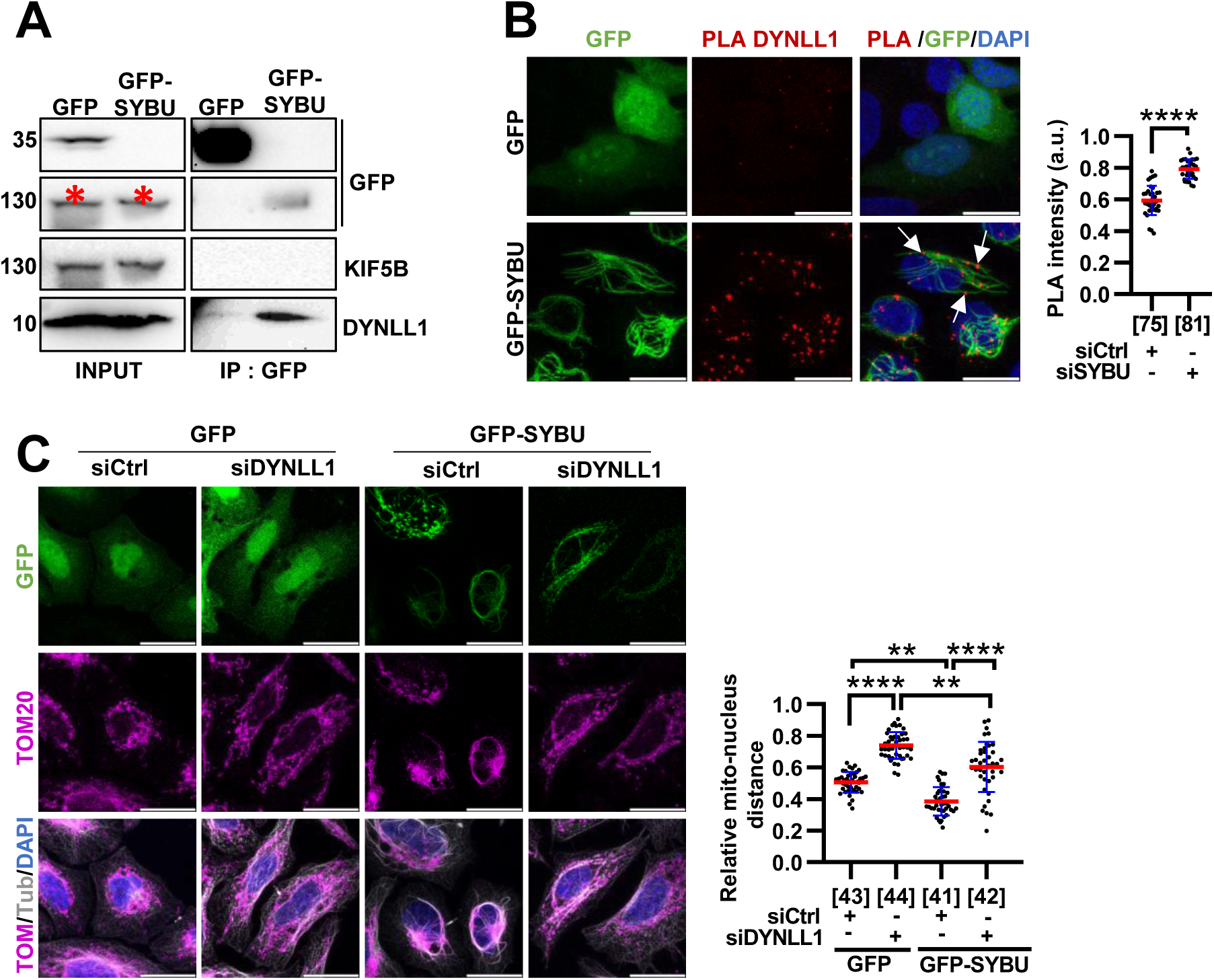
Syntabulin interacts with dynein light chain 1 *DYNLL1*. **(A)** Immunoprecipitation using anti-GFP antibodies in HeLa cells transfected with GFP or GFP-SYBU. Membranes were probed with anti-GFP, anti-KIF5B and anti-DYNLL1 antibodies. Molecular weight (KDa) is indicated on the left. Red asterisks indicate background signals from previous anti-KIF5B hybridization. **(B)** Proximity ligation assay (PLA) in HeLa cells transfected with GFP or GFP-SYBU, using anti-DYNLL1 and anti-GFP antibodies. Red puncta indicate proximity between GFP-SYBU and DYNLL1. Arrows show localization of PLA signals along microtubule bundles stained by GFP-SYBU. *Right:* Quantification of rolling circles PLA signal intensity. **(C)** Immunofluorescence of mitochondrial distribution in HeLa cells transfected with control siRNA (siCtrl) or siRNA targeting dynein light chain LC8-type 1 (siDYNLL1) and expressing GFP-SYBU or GFP as a control. *Right:* Quantification of relative distance between mitochondria and nucleus. **<u>Data information.</u>** Fluorescent staining shows PLA in red **(B)** and mitochondria in magenta and tubulin in grey **(C)**. Error bars represent mean ± SD from four **(B)** and three **(C)** independent experiments. Statistical analyses: **(B)** Mann-Whitney and **(C)** Kruskal-Wallis with Dunn’s *post hoc* tests. p**<0.01, p****<0.0001. Number of cells is indicated under brackets **(B,C)**. Scale bar = 10 µm.

At the functional level, DYNLL1 silencing mimicked *SYBU* depletion and favored mitochondria distribution at the cell periphery, confirming its role in mitochondria retrograde transport driven by cytosolic dynein (Fig. 4C). DYNLL1 depletion also reversed GFP-SYBU-induced perinuclear mitochondria localization (Fig. 4C). Similar resultats were obtained upon dynein inhibition by ciliobrevin D (Supplementary Fig. S4E), highlighting the essential role of dynein molecular motor in mitochondrial distribution controled by syntabulin. Collectively, these results indicate that syntabulin cooperates with dynein to regulate mitochondrial transport in cancer cells.

### Syntabulin opposes Miro1/KIF5B-mediated anterograde mitochondrial transport

Mitochondria peripheral positioning driven by DYNLL1 depletion was reversed by silencing the opposing molecular motor kinesin-1 (KIF5B) (Supplementary Fig. S5A-S5C). We thus tested whether loss of *SYBU* disturbs the balance between dynein and KIF5B to control mitochondria intracellular localisation. Silencing of KIF5B in *SYBU*-depleted cells indeed reversed the peripheral distribution of mitochondria as seen by super-resolution STORM microscopy (Fig. 5A), indicating that KIF5B mediates the anterograde mitochondrial transport caused by *SYBU* depletion. Expression of a constitutively active mutant of KIF5B (K560) phenocopied *SYBU* depletion in driving peripheral distribution of mitochondria in control cells (Fig. 5B). Conversely, expression of a motor-defective, immotile KIF5B mutant (rigor) prevented cortical mitochondrial distribution in *SYBU*-deficient cells (Fig. 5B), indicating that the motor activity of KIF5B is essential for anterograde mitochondrial transport in the absence of syntabulin.

**Figure 5.**
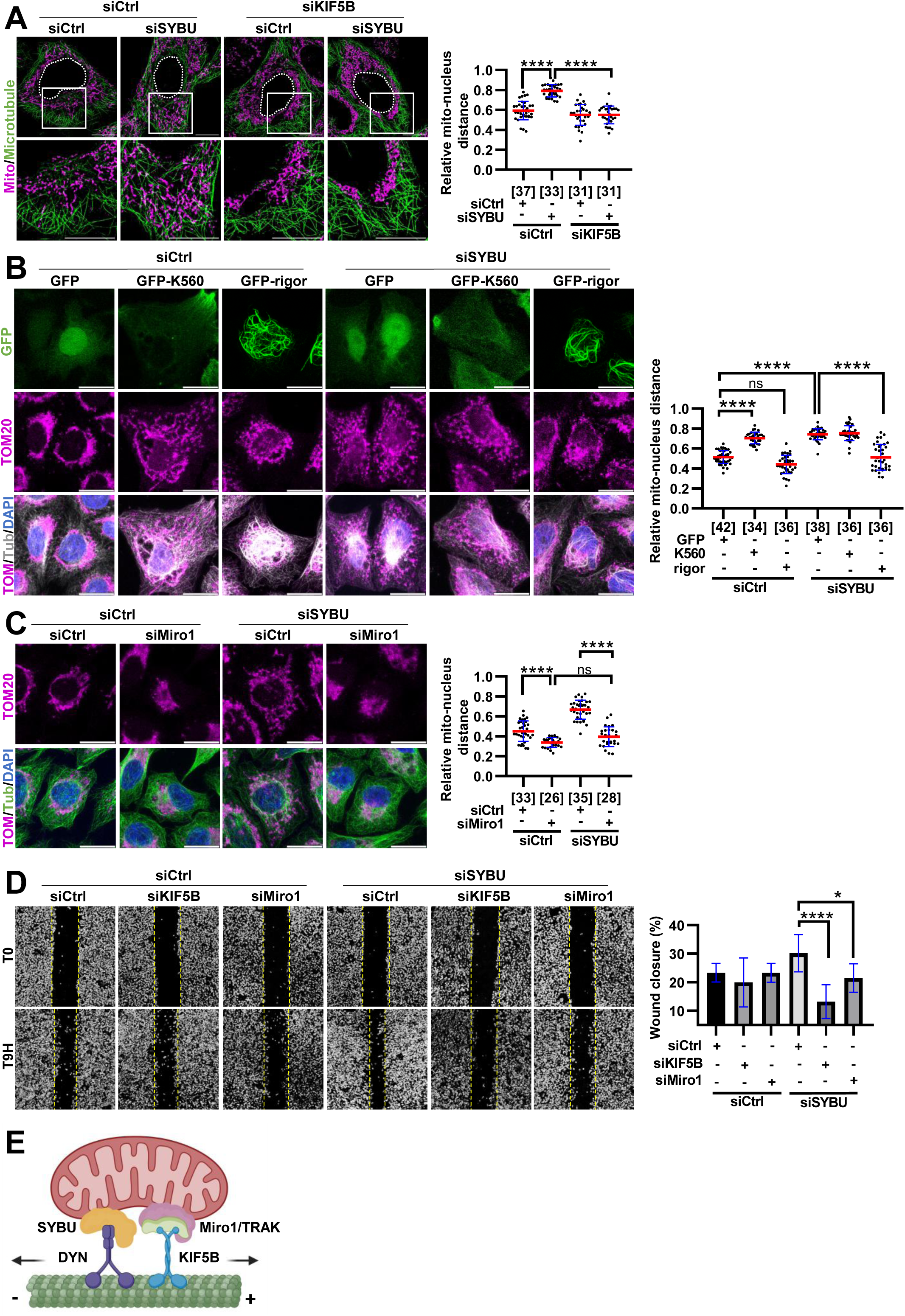
Syntabulin opposes Miro1/KIF5B-mediated anterograde mitochondrial transport. **(A)** Super resolution STORM microscopy of mitochondria distribution in resting HeLa cells transfected with siCtrl, siSYBU or siRNA targeting KIF5B (siKIF5B). *Right:* Quantification of relative distance between mitochondria and nucleus. **(B)** Immunofluorescence of mitochondria distribution in siCtrl or siSYBU-transfected HeLa cells expressing GFP, KIF5B-constitutively active GFP-K560 or immotile GFP-rigor mutant. *Right:* Quantification as (A). **(C)** Same as (B) with siRNA targeting Miro1 (siMiro1). *Right:* Quantification as (A). **(D)** Wound healing assays in siCtrl, siSYBU, siKIF5B or siMiro1-transfected D3H2LN cells. *Right:* Quantification of wound closure 9 hrs after scratching. **(E)** Proposed model of syntabulin/dynein (SYBU/DYN) complex opposing the anterograde transport of mitochondria mediated by the Miro1/TRAK/KIF5B machinery. Arrows indicate movement toward minus (-) or plus (+) end of microtubule. **<u>Data information.</u>** Fluorescent staining shows **(A-C)** mitochondria in magenta and microtubules in green; **(B)** tubulin in grey. Error bars represent mean ± SD from three **(A,B,D)** and two **(C)** independent experiments. **(D)** n=4 wells per experiment. Statistical analyses: **(A,C)** One-way ANOVA with Tukey’s *post hoc* and **(B,D)** Kruskal-Wallis with Dunn’s *post hoc* tests. p*<0.05, p****<0.0001. Number of cells is indicated under brackets. Scale bar = 10 µm.

Since KIF5B transports mitochondria *via* the mitochondrial adaptors Miro1/TRAK, we investigated whether Miro1 governs mitochondrial positioning in the absence of syntabulin. We found that Miro1 depletion prevents mitochondria peripheral distribution induced by *SYBU* depletion (Fig. 5C; Supplementary Fig. S5D), indicating that syntabulin opposes the effects of both Miro1 and KIF5B on mitochondrial transport. We then questioned the functional relevance of these findings and explored the consequence of depleting Miro1 and KIF5B on cell migration. We found that silencing either KIF5B or Miro1 in breast cancer cells reversed the pro-migratory phenotype caused by *SYBU* depletion (Fig. 5D), highlighting the relevance of Miro1/KIF5B-mediated mitochondrial distribution in the migratory properties of *SYBU*-deficient cancer cells.

Together, these results identify syntabulin as a new partner of dynein and a negative regulator of the Miro1/KIF5B pathway (Fig. 5E), acting to limit mitochondrial anterograde transport and consequently, breast cancer cell migration.

### *SYBU* depletion promotes microtubule deacetylation and damage

The movement of KIF5B along microtubules has been shown to create friction forces and local lattice damage that serve as entry points for the HDAC6 tubulin deacetylase into the lumen of the microtubule, with subsequent deacetylation of α-tubulin (Andreu-Carbó et al., 2024). This led us to investigate whether *SYBU* deficiency, by promoting KIF5B-mediated anterograde transport of mitochondria, also causes microtubule damage and deacetylation.

*SYBU* silencing in HeLa cells led to a significant decrease in microtubule acetylation (Fig. 6A, Supplementary Fig. S6A-S6C), which was reversed by GFP-SYBU expression (Fig. 6B). Similar results were obtained in CAL120 and D3H2LN breast cancer cells (Supplementary Fig. S6D-S6G). The reduction in microtubule acetylation observed upon *SYBU* depletion was reversed by silencing KIF5B (Fig. 6C). Expression of the constitutively active mutant (K560) of KIF5B mimicked the effects of *SYBU* depletion whereas the immotile KIF5B mutant (rigor) restored microtubule acetylation in the absence of syntabulin (Fig. 6D). Thus, the motor activity of KIF5B is essential for decreasing microtubule acetylation in *SYBU*-depleted cells. As for syntabulin, DYNLL1 silencing decreased microtubule acetylation and this phenotype was also rescued by KIF5B silencing (Supplementary Fig. S6H), further highlighting the opposite effects of dynein and KIF5B, and supporting our proposed model (Fig. 5E).

**Figure 6.**
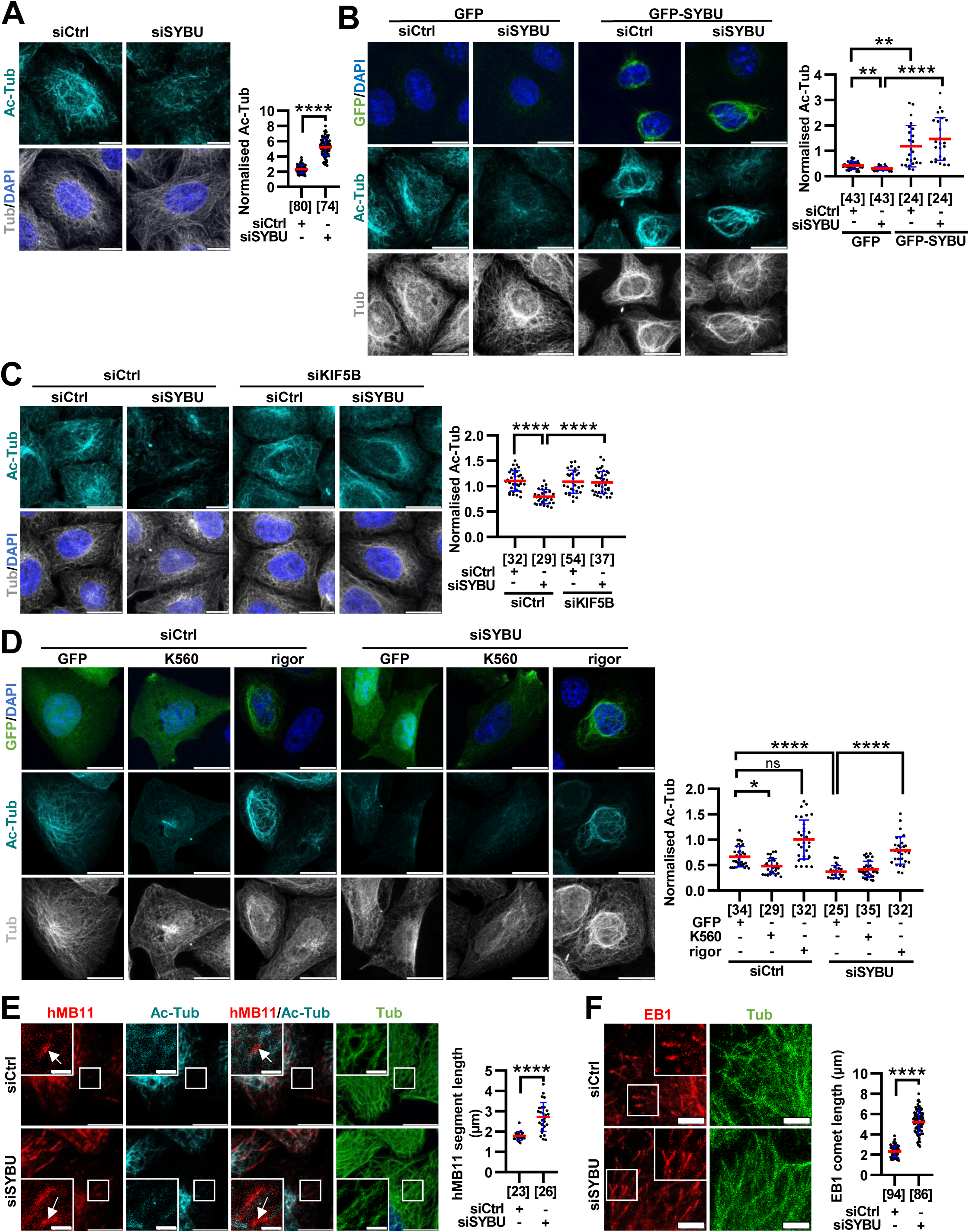
*SYBU* depletion decreases microtubule acetylation and causes microtubule damage. **(A)** Immunofluorescence of microtubule acetylation in siCtrl or siSYBU-transfected HeLa cells. *Right:* Quantification of acetylated tubulin (Ac-Tub) relative to tubulin (Tub) intensity. **(B)** Same as (A) after transfection with GFP or GFP-SYBU. *Right:* Quantification as (A). **(C)** Same as (A) in siCtrl, siSYBU or siKIF5B-transfected HeLa cells. *Right:* Quantification as (A). **(D)** Same as (A) after transfection with GFP or GFP-KIF5B mutants (K560 or rigor). *Right:* Quantification as (A). **(E)** Immunofluorescence of microtubule damage/repair sites in siCtrl or siSYBU-transfected HeLa cells stained with anti-hMB11 (recognizes GTP-tubulin). *Right:* Quantification of hMB11 segment length. **(F)** Same as (E) with anti-EB1 staining (recognizes growing microtubule ends). *Right:* Quantification of EB1 comet length. **<u>Data information</u>**. Fluorescent staining shows **(A-D)** Ac-Tub in cyan and Tub in grey; **(E,F)** hMB11 and EB1 in red and Tub in green. Error bars represent mean ± SD from three **(A,C,D,F)** and two **(E)** independent experiments. **(B)** Shown is one representative out of two experiments. Statistical analyses: **(A,E,F)** Mann-Whitney; **(B,D)** Kruskal-Wallis with Dunn’s *post hoc* and **(C)** One-way ANOVA with Tukey’s *post hoc* tests. p**<0.01, p****<0.0001. Number of cells is indicated under brackets. Scale bar = 10 µm **(A-D)**, 4 µm **(F)**, 20 µm **(E).** Zoom = 4 µm **(E)**.

In line with decreased microtubule acetylation, *SYBU* silencing also increases microtubule damage, revealed by staining with hMB11 antibodies (Dimitrov et al., 2008) (Fig. 6E). Furthermore, the size of EB1 comets, surrogate markers for growing plus ends which are other key entry sites for HDAC6, was also increased in *SYBU-*depleted cells (Fig. 6F). Collectively, these data suggest that in *SYBU*-deficient cells, enhanced anterograde mitochondrial transport driven by KIF5B contributes to microtubule damage and subsequent deacetylation.

### Microtubule deacetylation favors KIF5B-mediated anterograde mitochondrial transport

Microtubule acetylation by itself has been described as a general mechanism implicated in modulating kinesin-1 binding and motor activity. Here, we show that reducing microtubule acetylation by silencing either αTAT1 (the enzyme responsible for transferring acetyl-CoA to lysine 40 of α-tubulin) or ACSS1 and ACSS2 (two enzymes involved in acetyl-CoA production) leads to peripheral distribution of mitochondria, independently of syntabulin (Fig. 7A-7C; Supplementary Fig. S7A). Furthermore, expression of a non-acetylable α-tubulin mutant (K40A) mimicked *SYBU* depletion in promoting mitochondria positioning close to the periphery (Fig.7D; Supplementary Fig. S7B). These results support the notion that microtubule deacetylation by itself may drive peripheral localization of mitochondria.

**Figure 7.**
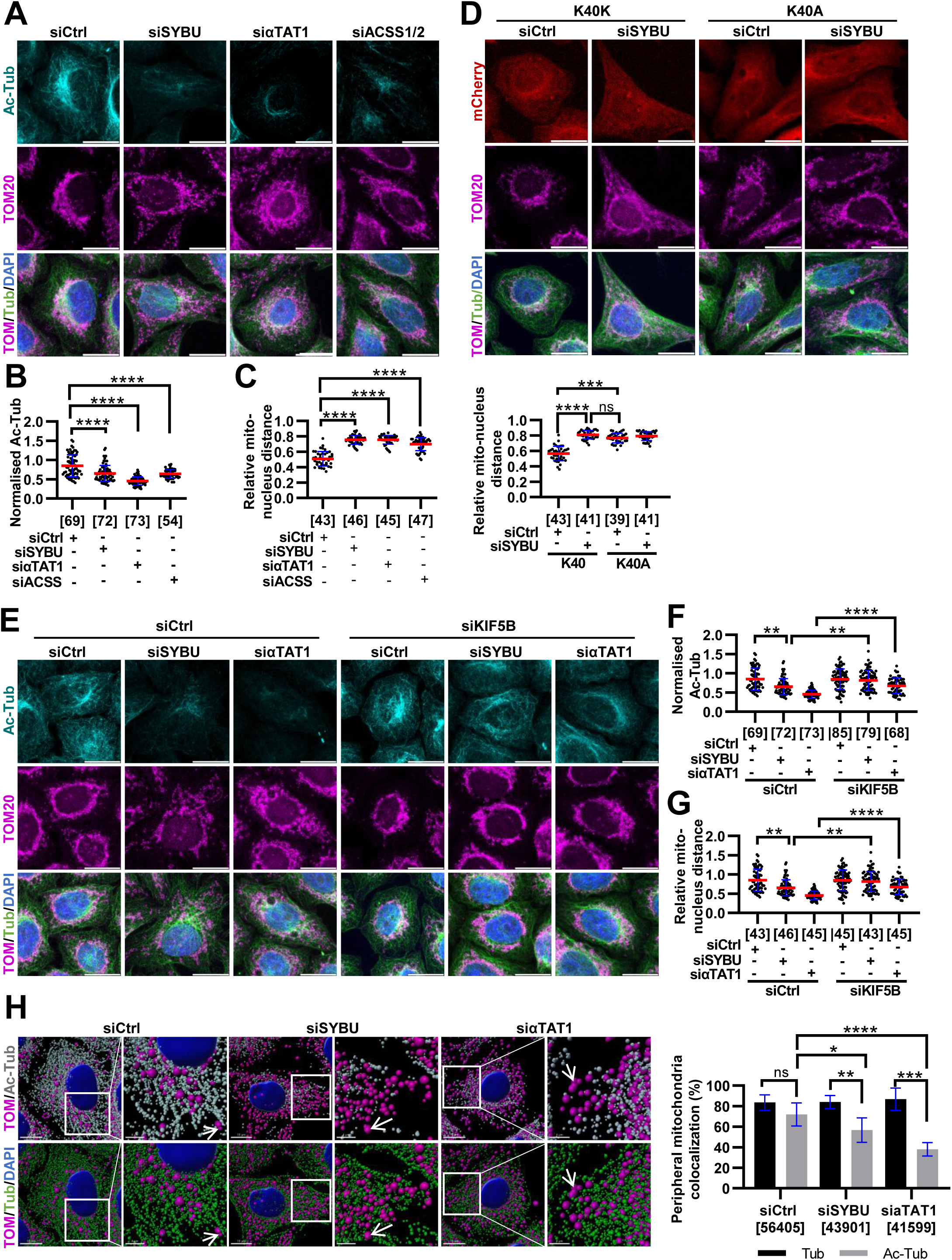
Microtubule deacetylation promotes anterograde mitochondrial transport. **(A)** Immunofluorescence of mitochondria distribution and microtubule acetylation in HeLa cells transfected with siCtrl, siSYBU, siαTAT1 (siRNA targeting αTAT1) or siACSS1/2 (siRNA targeting ACSS1 and ACSS2). **(B)** Quantification from (A) of Ac-Tub relative to total tubulin (Tub) intensity. **(C)** Quantification from (A) of relative distance between mitochondria and nucleus. **(D)** Immunofluorescence of mitochondria distribution in HeLa cells transfected with mCherry-K40K (wild-type α-tubulin) or mCherry-K40A (non-acetylable). *Bottom:* Quantification as (C). **(E)** Same as (A) in siCtrl, siSYBU, siαTAT1 or siKIF5B-transfected HeLa cells. **(F)** Quantification from (E) as (B). **(G)** Quantification from (E) as (C). **(H)** 3D-segmented images in siCtrl, siSYBU or siαTAT1-transfected HeLa cells. Arrows show mitochondria attached to non-acetylated microtubules. *Right:* Quantification of the percentage of peripheral mitochondria co-localized with microtubules (black) or acetylated microtubules (grey). **<u>Data information</u>**. Fluorescent staining shows **(A,E,H)** mitochondria in magenta and tubulin in green; **(A,E)** Ac-Tub in cyan; **(H)** Ac-Tub in grey. Error bars represent mean ± SD from three **(A-G)** or two **(H)** independent experiments. Statistical analyses: **(B-G)** Kruskal-Wallis with Dunn’s *post hoc* and **(H)** Two-way ANOVA with Sidak’s multiple comparisons tests. p*<0.05, p**<0.01, p***<0.001, p****<0.0001. Number of cells is indicated under brackets **(B-G)**. For **(H)** under brackets is the number of mitochondria quantified in 184 siCtrl-, 167 siSYBU- and 128 siαTAT1-transfected cells. Scale bar = 10 µm. Zoom = 4 µm **(H)**.

Notably, KIF5B silencing reversed both mitochondrial mislocalization and microtubule deacetylation caused by αTAT1 silencing (Fig. 7E-7G), indicating that the anterograde mitochondrial transport occurring upon reduced microtubule acetylation is promoted by KIF5B. These data raised the question of whether mitochondria reaching the cell periphery in *SYBU* deficient cells are attached to acetylated or deacetylated microtubules. We performed image segmentation analysis, which revealed that mitochondria positioned at the vicinity of the periphery are co-localized with microtubules (Fig. 7H). In control cells, they are equally co-localized with both acetylated and non-acetylated microtubules. Interestingly, in *SYBU*- and αTAT1-deficient cells, where most microtubules are deacetylated, mitochondria distributed at the cell periphery are predominantly co-localized with non-acetylated microtubules (Fig. 7H). These findings suggest that when microtubule acetylation is limited, mitochondria are transported by KIF5B toward the plus ends of deacetylated microtubules.

Collectively, these data identify KIF5B kinesin as a motor protein that not only promotes microtubule deacetylation but also utilizes deacetylated microtubules to mediate mitochondrial transport.

### *SYBU*-depleted cells are vulnerable to HDAC6 inhibition

We reasoned that preventing microtubule deacetylation using HDAC6 inhibitors may restore mitochondrial mislocalization in *SYBU*-depleted cells. Treatment with HDAC6 inhibitor ACY-1215 (Ricolinostat) indeed restored the defects in both microtubule acetylation and mitochondrial distribution induced by *SYBU* depletion (Fig. 8A-8C). Similar results were obtained with HDAC6 inhibitor Tubacin and pan-HDAC inhibitor Vorinostat (SAHA) that rescued the *SYBU*-deficient phenotypes (Supplementary Fig. S8). Notably, ACY-1215 reversed the pro-migratory phenotype induced by *SYBU*-depletion (Fig. 8D), further highlighting the functional impact of microtubule acetylation on *SYBU*-deficient cancer cell migration, and positioning HDAC6 inhibitors as interesting anti-cancer drugs.

**Figure 8.**
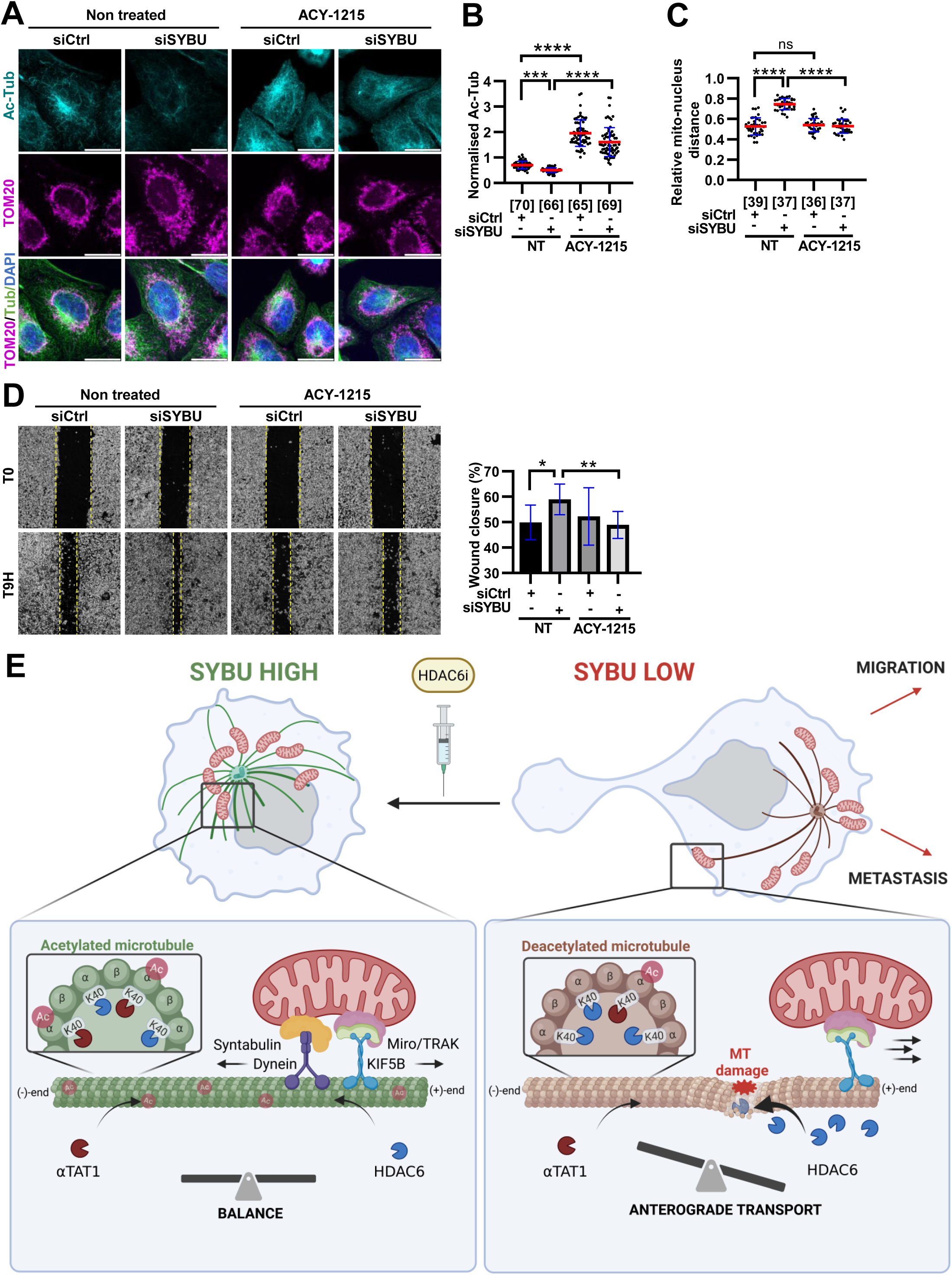
HDAC6 inhibition rescues *SYBU* silencing phenotypes. **(A)** Immunofluorescence of mitochondria distribution and microtubule acetylation in siCtrl or siSYBU-transfected HeLa cells, treated or not with 50 nM Ricolinostat (ACY-1215) during 4 hrs. **(B)** Quantification from (A) of Ac-Tub relative to Tub intensity. **(C)** Quantification from (A) of relative distance between mitochondria and nucleus. **(D)** Wound healing assays in siCtrl or siSYBU-transfected D3H2LN cells. Cells were pre-treated with 250 nM Ricolinostat (ACY-1215) for 1 hr, prior to wound healing assay for 9 hrs in the presence of Ricolinostat. *Right:* Quantification of wound closure. **(E)** Schematic representation of our proposed model. *Left:* In control condition (SYBU high), syntabulin and dynein form a molecular complex that opposes the Miro1/TRAK/KIF5B mediated anterograde transport of mitochondria. Microtubule acetylation levels are dynamically regulated by a balance between αTAT1 tubulin acetyltransferase and HDAC6 tubulin deacetylase. *Right:* in SYBU-deficient cells (SYBU-low), the balance is disrupted and KIF5B transports mitochondria to the cell cortex, causing microtubule damage (red star) and increased entry of HDAC6 in the lumen. The resulting microtubule deacetylation, in turn, facilitates KIF5B-mediated movement of mitochondria to the microtubule plus (+) end, with subsequent increase in cell migration and potentially breast cancer metastasis. HDAC6 inhibitor rescues *SYBU*-deficiency. **<u>Data information</u>**. Fluorescent staining shows **(A)** mitochondria in magenta, Ac-Tub in cyan and Tub in green. **(B-D)** Error bars represent mean ± SD from three independent experiments. Statistical analyses: **(B)** Kruskal-Wallis with Dunn’s *post hoc* test and **(C)** One-way ANOVA with Tukey’s *post hoc* test. p*<0.05, p**<0.01, p***<0.001, p****<0.0001. Scale bar = 10 µm.

## Discussion

Mitochondrial positioning is a critical determinant of cell fate, notably influencing cancer cell migration and invasion. Here we identify syntabulin, a mitochondrial outer membrane protein initially characterized in neurons, as a key regulator of mitochondrial transport in cancer cells. We show that *SYBU*, the gene encoding syntabulin, is a novel candidate prognostic biomarker that is downregulated in metastatic breast cancer. Loss of syntabulin enhances breast cancer cell migration and promotes anterograde transport of mitochondria toward the cell cortex, where they localize in close proximity to focal adhesions.

Mechanistically, syntabulin interacts with dynein along microtubule tracks and limits the anterograde mitochondrial transport mediated by the Miro1/KIF5B complex, thereby limiting cancer cell migration. In *SYBU*-depleted cells, the balance between dynein and KIF5B activities is shifted and mitochondria achieve faster and more persistent movements, covering longer anterograde than retrograde distances, which ultimately leads to their peripheral distribution. By promoting KIF5B movement along microtubules, *SYBU* depletion generates microtubule damage that serves as an entry point for HDAC6 deacetylase. Consistent with previous studies (Andreu-Carbó et al., 2024), we show that KIF5B motility in *SYBU*-depleted cells ultimately drives microtubule deacetylation. In turn, reduced microtubule acetylation, achieved either through *SYBU* depletion or by silencing αTAT1 or ACSS1/2, promotes mitochondrial transport to the cell periphery *via* KIF5B.

Our findings support a model (Fig. 8E) in which tight control of both microtubule acetylation and a dynein/KIF5B balance is critical for mitochondria positioning within cancer cells. In *SYBU*-depleted cancer cells, enhanced KIF5B motility provokes microtubule deacetylation which in turn facilitates KIF5B movement, establishing a positive feedback loop that reinforces peripheral mitochondrial localization. Importantly, treatment with the HDAC6 inhibitor ACY-1215 restored proper mitochondrial distribution and prevented the enhanced migratory behavior of *SYBU*-deficient cells.

We show here that KIF5B orchestrates the interplay between microtubule acetylation and mitochondria positioning in cancer cells, acting both to promote microtubule deacetylation and to utilize deacetylated microtubules for mitochondrial anterograde transport. These data provide new insights into the reciprocal regulation between microtubule acetylation and KIF5B-mediated anterograde transport (Reed et al., 2006; Tas et al., 2017; Ravindran et al., 2017; Monteiro et al., 2023), that remains an active field of investigation (Verhey and Ohi, 2023).

They also carry important clinical implications for breast cancer management. Our findings indicate that *SYBU*-deficient breast tumors, which account for approximately 50% of invasive breast tumors and 60% of metastatic cases, may be particularly susceptible to therapies targeting microtubule deacetylation. The selective HDAC6 inhibitor ricolinostat is an orally available compound currently under clinical evaluation in patients with metastatic breast cancer (Zeleke et al., 2023). Recent studies highlight the therapeutic potential of an emerging class of selective HDAC6 inhibitors to improve outcomes in breast cancer, with particular relevance to TNBC patients (Huang et al., 2016, 2024). The identification of syntabulin, a mitochondrial outer membrane protein, as a novel regulator of microtubule acetylation provides a mechanistic framework to better understand how alterations in the mitochondria-microtubule cross-talk contribute to metastatic dissemination. Importantly, it also offers new preclinical and translational opportunities to evaluate the therapeutic value of targeting microtubule deacetylation in a personalized way for *SYBU*-low breast cancer patients, a subgroup characterized by poor clinical outcome.

## Material and Methods

### Breast cancer patients cohorts

Microarray analyses were performed on tumor biopsies from the French REMAGUS-02 (R02) cohort, a randomized phase II neoadjuvant clinical trial including 115 breast cancer patients. The clinical and pathological characteristics of this cohort have been described previously (Rodrigues-Ferreira et al., 2019).

Expression levels of the SYBU/GOLSYN gene in normal, tumor, and metastatic breast tissues were obtained from publicly available transcriptomic datasets comprising a total of 7,893 breast cancer patients using the TNMplotdatabase (Bartha and Győrffy, 2021). Patient survival analyses were performed using the KMplot online database (Győrffy, 2021). For each analysis, the best probe set cutoff was applied.

### Cell lines and culture conditions

Experiments were done on metastatic TNBC cell lines: MDA-MB-231-luc-D3H2LN (Jenkins et al., 2003) (Rodrigues-Ferreira et al., 2012), as well as HeLa (cervical carcinoma cell line). Cells were routinely authenticated by morphologic observation and periodically tested for absence of mycoplasma contamination using Venor® GeM Advance Kit (Venor®GeM Advance kit). Cells were grown in complete medium (DMEM Dulbecco’s Modified Eagle Medium + 10% FBS (Fœtal Bovine Serum) and incubated in 5% CO_2_ at 37°C. Cell seeding conditions are specified in the Supplementary Materials section. Ricolinostat treatment (MedChemExpress, HY-16026-1ML; 0,5 nM to 5 µM for 4 to 9 hrs) was performed in complete medium, after cell transfection. Treatment duration and concentration were optimized based on preliminary dose-response experiments.

### siRNA and plasmid transfection

Cells were transfected at ~80% confluence with siRNAs (20 nM) using Lipofectamine 2000 (Thermo Fisher Scientific), diluted in Opti:MEM according to the manufacturer’s instructions. Non-targeting siRNA (siCtrl) was used as a negative control (Lifetech ambion, ref 4390844). siRNA sequences purchased from Dharmacon (Chicago, IL, USA) are provided in the Supplementary Materials and Methods section.

For plasmid transfection, cells were transfected at ~60% confluence using Lipofectamine 2000 (Thermo Fisher Scientific) with 1-2 µg of plasmid DNA per 6-well plate. For rescue experiments with GFP-SYBU, the construct was made resistant to siRNA by using QuikChange® II XL Site-Directed Mutagenesis Kit and oligonucleotide sequence: 5’-CTGGAAGGTACATGTCCTGTGGAGAAAATCATGGTGGTC-3’.

GFP-K560 and GFP-rigor plasmids were described elsewhere (Andreu-Carbó et al., 2024), and mCherry-K40 and mCherry-K40A plasmids here (Dompierre et al., 2007).

### Cell migration and invasion

For scratch and Transwell assays, technique was previously described (Rodrigues-Ferreira et al., 2012). Briefly, scratch assay was performed on cell monolayer reaching full confluence and scratched using a sterile pipette tip. Images of the wound area were taken at time 0 and at different time points using a phase-contrast microscope (Invitrogen™ EVOS XL Core System). Wound closure was quantified manually using ImageJ by measuring the scratch area over time. *% Wound closure = ((Area_T(₀)_ - Area_T(X)_) / Area_T(₀)_) × 100*.

For transwell migration assay in Boyden Chamber, cells in serum-free medium were seeded into the upper chamber of transwell inserts with 8 µm pore polycarbonate membranes (Corning® Costar, 24-well format). The lower chamber was filled with complete growth medium containing 10% FBS as a chemoattractant. In parallel, cells were incubated in wells without a filter to determine the total number of seeded cells. After 6 hrs of incubation, cells that migrated on the lower surface were fixed, stained with crystal violet, and imaged (phase-contrast microscope). Cells were quantified by manually counting in ImageJ.

For spheroids invasion, cells were seeded into 96-well plates treated with PolyHema 1X to reduce cell attachment, and briefly centrifuged to promote aggregation. Matrigel (Geltrex) 200 µg/ml was carefully added to polymerize around cells. Spheroids were allowed to form over 48 hrs. A denser Matrigel matrix (1 mg/mL) was subsequently overlaid. Invasion was monitored over 24 hrs using phase-contrast microscope (Incucyte® Sartorius) and spheroids invasion was calculated as: *Area_T(₀)_ - Area_T(X)_*.

### Immunofluorescence staining

For mitochondria and microtubules co-staining, cells were fixed with 4% paraformaldehyde (PFA) heated to 37°C for 15 minutes at room temperature, followed by permeabilization with 0.1% Triton X-100. Primary antibodies were incubated 1 hr at 37°C in blocking solution PBS 1X BSA 1%: anti-TOM20 clone D8T4N (Cell Signaling, 42406, rabbit) or anti-TOM20 F-10 (Santa-Cruz, sc-17764, mouse); anti-tubulin (Abcam, ab6160, rat); anti-acetyl-tubulin (Sigma, T6793-100, mouse); anti-DYNLL1 (Abcam, ab51603, rabbit); anti-KIF5B (Abcam, ab167429, rabbit); anti-paxillin (Cell Signaling, E6R6Z, rabbit); anti-GFP [9F9.F9] (Abcam, ab1218, mouse). For microtubule staining, cells were fixed in cold methanol (−20°C) for 5 minutes and stained with anti-EB1 [KT51] (ThermoFisher, MA1-72531, rat). Secondary antibodies, purchased from Jackson Immunoresearch, were incubated 45 minutes at 37°C in blocking solution PBS 1X BSA 1%. Nuclei were counterstained with DAPI (Invitrogen), and coverslips were mounted using FluorSave reagent (Millipore).

For anti-hMB11 staining (human antibody against GTP-MTs; Institut Curie, Paris, France), method has been described previously (Dimitrov et al., 2008). Briefly, free-tubulin was extracted in 0.1% Triton X-100 in warm PEM buffer. Cells were then incubated for 15 minutes with anti-hMB11 antibody in PEM-1% BSA, followed by washes in PEM. Fixation was performed for 5 minutes in methanol at −20°C. Cells were then incubated for 15 minutes at room temperature with primary then secondary antibodies.

### Immunofluorescence quantifications

To quantify the relative distance between mitochondria and nucleus, we selected the mitochondria distributed nearest to the cell membrane and performed the measurements as illustrated in Supplementary Fig. S3C. For each mitochondria, we manually measured its distance to the cell periphery (d1) and the distance from the periphery to the nucleus (d2), using the measurement tool in LAS X software. The relative mito-nucleus distance was defined as: *1-(d1/d2).* In each cell, measurements were performed on 8 to 15 mitochondria, by radiating around the cell. The resulting values were averaged to obtain a mean relative distance per cell. Values are ranging from 0.0 (nucleus) to 1.0 (cell periphery).

For transfected cells, plasmid expression intensity was quantified for each condition. Cells were stratified by median intensity, and only the low-expressing population was analyzed.

To evaluate the distance between focal adhesions and mitochondria, the closest distance between the mitochondria and the focal adhesion were counted.

The ratio of acetylated tubulin to total tubulin was calculated as a measure of the overall acetylated tubulin staining intensity relative to total tubulin.

Finally, three-dimensional (3D) image segmentation and analysis were performed using Imaris 9 (Bitplane, Oxford Instruments). Individual mitochondria, total tubulin, acetylated tubulin and nuclei were segmented as individual objects. Segmentation parameters were optimized to accurately define object boundaries based on intensity thresholds and morphological criteria. Following automated segmentation, 3D renderings were visually inspected for accuracy and ensure proper object identification, following batch processing. Quantitative measurements, including object-based colocalization and spatial metrics, were extracted using Imaris’ built-in statistics tools. Specifically, colocalization was defined based on the Euclidean distance between segmented objects: mitochondria were considered colocalized with tubulin (total or acetylated) when the distance between their surfaces was less than 0.5 µm, and not colocalized when the distance exceeded this threshold. To analyze mitochondrial spatial distribution within the cell, the distance from the nucleus to each mitochondrial object was calculated. A distance threshold of 7 µm from the nuclear surface was used to distinguish between perinuclear mitochondria (< 7 µm) and peripheral mitochondria (> 7 µm). All relevant spatial and colocalization data were exported from Imaris for downstream quantitative analysis.

### Super-resolution imaging (SMLM Single Molecule Localization Microscopy)

Super-resolution imaging was performed using direct STORM (Stochastic Optical Reconstruction Microscopy) microscopy, with the ready-to-use Abbelight Smart Staining Kit Tubulin-Mitochondria (Abbelight, France). Cells were fixed, blocked, and permeabilized according to the manufacturer’s protocol. Immunostaining was performed with tubulin AF647 and mitochondria CF680. Samples were mounted in Abbelight imaging buffer, ensuring fluorophore photoswitching. Imaging was performed using a Nikon TiE2 inverted microscope with a 100X 1.49 NA objective, equipped with the SAFe module in HiLo illumination mode (Abbelight). To achieve simultaneous multicolor imaging, spectral demixing was applied, based on spectral properties. A minimum of 20,000-50,000 frames were acquired. Localization and reconstruction into 2D were carried out with Abbelight’s NEO software. Quantification of mitochondrial distribution was performed following the same procedure as used for standard immunofluorescence analyses.

### Time-lapse videomicroscopy of mitochondria trafficking

GFP-D3H2LN cells were incubated with MitoTracker Deep Red (Invitrogen M22426) 150 nM 1 hr prior to imaging. Time-lapse imaging was performed at 37°C in a humidified chamber with 5% CO₂. Mitochondrial movements were monitored using a spinning disk microscope (Nikon Eclipse Ti2) with 63X oil immersion objective. TIRF illumination was adjusted to selectively excite the basal region of the cell. Images were acquired every 2,5 seconds for 5 minutes, 30 minutes post-seeding, using minimal laser power to limit phototoxicity. Focus stability was maintained with Nikon Perfect Focus System. Mitochondrial trajectories were obtained by manual tracking with the TrackMate plugin in Fiji. The resulting (x,y) coordinates were exported and used for quantitative analyses. Mitochondrial transport velocity (µm/s) is defined as the distance traveled divided by time (number of frames x 2.5 seconds). Distance traveled by mitochondria (µm) corresponds to the sum of the cumulative displacements between each step, calculated as: 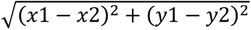; where (x_1_, y_1_) and (x₂, y₂) are the coordinates of two consecutive points. Persistence is defined as the ratio between the Euclidean distance and the total distance travelled by the mitochondria.The Euclidean distance is calculated as: 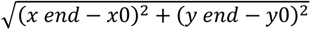; where *(x_0_, y_0_)* are the coordinates of the first point and *(x_end_, y_end_)* those of the last point.

### Proximity Ligation Assay (PLA)

Protein-protein interactions were analyzed using the Duolink® In Situ Red Starter Kit (Sigma-Aldrich) as described (Velot et al., 2015). Briefly, cells were fixed, permeabilized and blocked with Duolink blocking solution. Primary antibodies anti-GFP (Abcam, ab1218) and anti-DYNLL1 (Abcam, ab51603, rabbit) were incubated overnight at 4°C. Specific PLA probes conjugated to oligonucleotides were applied, followed by ligation and rolling circle amplification. Coverslips were mounted with Duolink mounting medium. Images were acquired using the confocal microscope Leica SP8. PLA signals were quantified as fluorescent puncta per cell using the Leica Application Suite X software. PLA intensity signals per region of interest (ROI) were quantified with the « polyline » tool.

### Co-immunoprecipitation

Cells were lysed in buffer, supplemented with protease and phosphatase inhibitors, as described (Nehlig et al., 2020). Lysates from GFP control vector or GFP-SYBU were centrifuged at 13.000 rpm for 15 minutes at 4 °C. To reduce nonspecific binding, lysates were incubated with non-conjugated magnetic beads (Chromotek) for 30 minutes at 4 °C under gentle rotation. Beads were then removed by magnetic separation, and the clarified lysate was collected. Supernatants were incubated for 2 hours at 4 °C with GFP-Trap magnetic beads (Chromotek), pre-equilibrated in lysis buffer. Beads were washed and bound proteins were eluted in Laemmli buffer. Eluted proteins and input lysates (1/3 of total lysate) were separated by SDS-PAGE and transferred onto PVDF membranes. Western blotting was performed using antibodies against DYNLL1, KIF5B and GFP. The western-blot was then performed as described in the Supplementary Materials and Methods section.

### Statistical analysis

Graphs were edited on the Graph Pad Prism® software (version 8). Data from TNMplot are compared with Dunn’Test. Data were first tested for normality (Anderson-Darling, D’Agostino & Pearson test, Shapiro-Wilk and Kolmogorov-Smirnov tests). For two groups, parametric data were analyzed using unpaired Student’s t-tests. In case of non-normally distributed data, non-parametric Mann-Whitney U tests were used. For more than two groups, parametric data were analyzed using ordinary one-way ANOVA followed by Tukey’s multiple comparisons. In case of non-normally distributed data, Kruskal-Wallis test was done. A difference between two conditions was considered as significative if p-value * < 0.05, ** < 0,01, *** < 0,001 and **** < 0.0001. ns = not significant.

## Acknowledgments

We wish to thank Tudor Manoliu (Université Paris-Saclay, INSERM US23 / CNRS UAR 3655, Imaging and Cytometry core facility PFIC, Gustave Roussy Institute, Villejuif, France) for excellent imaging expertise, Dr. Julien Courchet and Dr Martijn Kerkhofs (CNRS, Inserm, UMR5261, U1315, Institut NeuroMyoGène, 69008 Lyon, France) for helpful discussion and technological support. We also thank Paul Barthélémy (Abbelight) and Dr Daniel Perdiz (Paris-Saclay University) for providing reagents and constructive discussions throughout the project, and Auriane Debaumarché (Gustave Roussy Institute) for technical assistance. We are grateful to Dr Maria Haykal (Institute of Cellular Biochemistry and Genetics, University of Bordeaux) for valuable technical assistance and helpful discussions throughout this project.

