## Supplementary data for "Microtubule deacetylation drives kinesin-1 mediated mitochondrial transport accelerating breast cancer cell migration"

#### Supplementary Material and Methods

##### Breast cancer patients cohorts

The Curie cohort, the Gustave Roussy cohort (IGR), and the R04 cohort including 130 primary invasive ductal breast carcinomas, 177 invasive breast carcinomas and 142 breast cancer, respectively, were described elsewhere (Rodrigues-Ferreira et al., 2020; Molina et al., 2013; Rodrigues-Ferreira et al., 2019).

##### Cell lines and culture conditions

Experiments were also done on metastatic TNBC cell lines: MDA-MB-231 and CAL120. Cells were grown in complete medium (DMEM Dulbecco's Modified Eagle Medium + 10% FBS (Fœtal Bovine Serum) and incubated in 5% $CO_2$ at 37°C.

Drug treatments were performed with : Ciliobrevin D (MedChemExpress, HY-122632; 25 µM 1 hr); Tubacin (MedChemExpress, HY-16026-1ML; 2 µM 4 hrs); Vorinostat (MedChemExpress, HY-10221; 2 µM 4 hrs).

##### Cell seeding conditions

For fixed-cell imaging, cells were seeded on 1.5 coverslip glass (Ibidi).

For live-cell imaging, cells were seeded into µ-Slide 8 Well ibiTreat chambers (Ibidi).

For micropatterning experiments, cells were seeded on 1.5 coverslip glass featuring crossbow-shaped adhesive motifs. Micropatterns were generated using deep UV lithography on PEG-coated surfaces, creating protein-repellent backgrounds surrounding fibronectin-coated adhesive areas. Crossbow patterns coated with fibronectin (10 µg/mL) and cells were incubated for 15-30 minutes. Non-adherent cells were gently washed away.

##### siRNA sequences for transfection

*SYBU* (GUACAUGUCUUGCGGUGAA),

*KIF5B* (GAACUGGCAUGAUAGAUGA),

*RHOT1* (UGUGGAGUGUUCAGCGAAA),

*DYNLL1* (GUUCAAAUCUGGUUAAAAG; GAAGGACAUUGGGCUCAU; GUACUAGUUUGUCGUGGUU; CAGCCUAAAUUCCAAALAC).

Knockdown efficiency was confirmed by qRT-PCR/immunoblotting 48-72 hrs post-transfection. Plasmid transfection was validated by immunoblotting/immunofluorescence.

##### Spheroid growth assay

For spheroids growth, cells were seeded into 96-well plates treated with PolyHema 1X to reduce cell attachment, and briefly centrifuged to promote aggregation. Matrigel (Geltrex) 200 µg/ml was carefully added to polymerize around cells. Spheroids were allowed to form over 48 hrs. Growth was monitored over 9 days using phase-contrast microscope. Spheroids growth was quantified by manually measuring spheroid area using ImageJ software, and calculated as $Area_{T(0)} / Area_{T(X)}$.

**Immunofluorescence acquisition**

Images were acquired on the confocal microscope Leica SP8 using an immersion oil objective X63. Each image has a resolution of 12 bit in a frame of 2048x2048. To capture the entire cell, the Z-stack was made on 10 steps of 0.8 µm. The software used was the Leica Application Suite X (v3.5.7.23225). Representative images were then extracted (maximum intensity).

**Western blotting**

Cells were lysed in RIPA buffer supplemented with protease and phosphatase inhibitors. Syntabulin antibody has been revealed after sonication. Protein concentrations were determined using the BCA assay (Pierce). Equal amounts of protein were denatured in Laemmli buffer and separated on SDS-PAGE 10% gels. Proteins were transferred onto PVDF membranes (Thermo Fisher, 7-minute program, iBlot Transfer Stack). Membranes were blocked in 5% non-fat dry milk in Tris-buffered saline with 0.1% Tween-20. Then, incubated overnight at 4 °C with primary antibodies : anti-syntabulin (Abcam, ab138528, rabbit) ; anti-tubulin (Abcam, ab6160, rat); anti-acetyl-tubulin (Sigma, T6793-100, mouse); anti-vinculin (Sigma, V9264, mouse); anti-DYNLL1 (Abcam, ab51603, rabbit); anti-KIF5B (Abcam, ab167429, rabbit). After incubation with HRP-conjugated secondary antibodies, signal was visualized using enhanced chemiluminescence (Clarity Western ECL substrate) and imaged on a ChemiDoc MP System (Bio-Rad). Band intensities were quantified using Image Lab software and normalized to loading controls.

**Quantitative PCR (qPCR)**

RNA extraction and reverse transcription were performed as previously described (Rodrigues-Ferreira et al., 2019). In brief, total RNA was extracted using TRIzol Reagent (Invitrogen) following phase separation with chloroform and precipitation with isopropanol. RNA pellets were washed with 75% ethanol, resuspended in nuclease-free water, and quantified using NanoDrop spectrophotometer. For reverse transcription, 1 µg of RNA was incubated with 10 mM oligo(dT) primers at 65 °C for 10 minutes, then cooled on ice. First-strand synthesis was performed using SuperScript® II reverse transcriptase (Thermo Fisher), DTT, RNaseOUT, and dNTPs at 37 °C for 1 hour, followed by inactivation at 70 °C for 5 minutes. qPCR was conducted using 2.5 ng/µL cDNA, gene-specific primers (10 µM), and Luminaris HiGreen

Master Mix in MicroAmp® plates, on a ViiA™ 7 system with SYBR Green detection and a standard $2^{-\Delta\Delta Ct}$ analysis normalized to a housekeeping gene.

**Primers for quantitative PCR (qPCR)**

*SYBU* Forward (F) : AAT CTG AGC GCC GAC TCC ATG A ;
*SYBU* Reverse (R) : CTT TCC TGG CTT CTT TGA GTG CC ;
*KIF5B* (F) : GAG TTA GCA GCA TGT CAG CTT CG;
*KIF5B* (R) : GCA TCG ACA GAT TCC TCC AAC TG ;
*DYNLL1* (F) : GCT ACT CAG GCG CTG GAG AAA T ;
*DYNLL1* (R) : GTG TCA CAT AAC TAC CGA AGT TCC ;
*MIRO1* (F) : GAC AAA GAC AGC AGG CTG CCT T ;
*MIRO1* (R) : TCG CTG AAC ACT CCA CAC AGG T ;
*RPL13* (F) : CTC AAG GTG TTT GAC GGC ATC C ;
*RPL13* (R) : TAC TTC CAG CCA ACC TCG TGA G.

**A**

| | | | |
|---|---|---|---|
| | Breast samples | | |
| **SYBU Expression** | Normal (242) | Tumor (7569) | Metastatic (82) |
| min | 22 | 1 | 30 |
| Q1 | **767** | 314 | 177 |
| med | 1346 | 676 | 449, 5 |
| Q3 | 2582 | 1175 | 987 |
| max | 8975 | 12505 | 3080 |

**B**

| | | | |
|---|---|---|---|
| | | Breast cancer samples | |
| | | Tumor (7569) | Metastatic (82) |
| | number of Samples with SYBU expression < Q1 (767) | 4189 | 49 |
| | **%** | **55,3** | **59,8** |

#### Table S1. SYBU gene expression in normal, tumor and metastatic breast samples

Analysis of *SYBU* gene expression in normal (n=242), tumor (n=7569), and metastatic breast samples (n=82), obtained from the TNMplot database. **(A)** Shown are minimum (min), quartiles (Q1, Q3), median (med), and maximum (max) of *SYBU* expression values for the three groups. **(B)** Percentage of tumor (55.3%) and metastatic (59.8%) samples with *SYBU* expression below the first quartile (Q1) of normal samples (Q1=767).

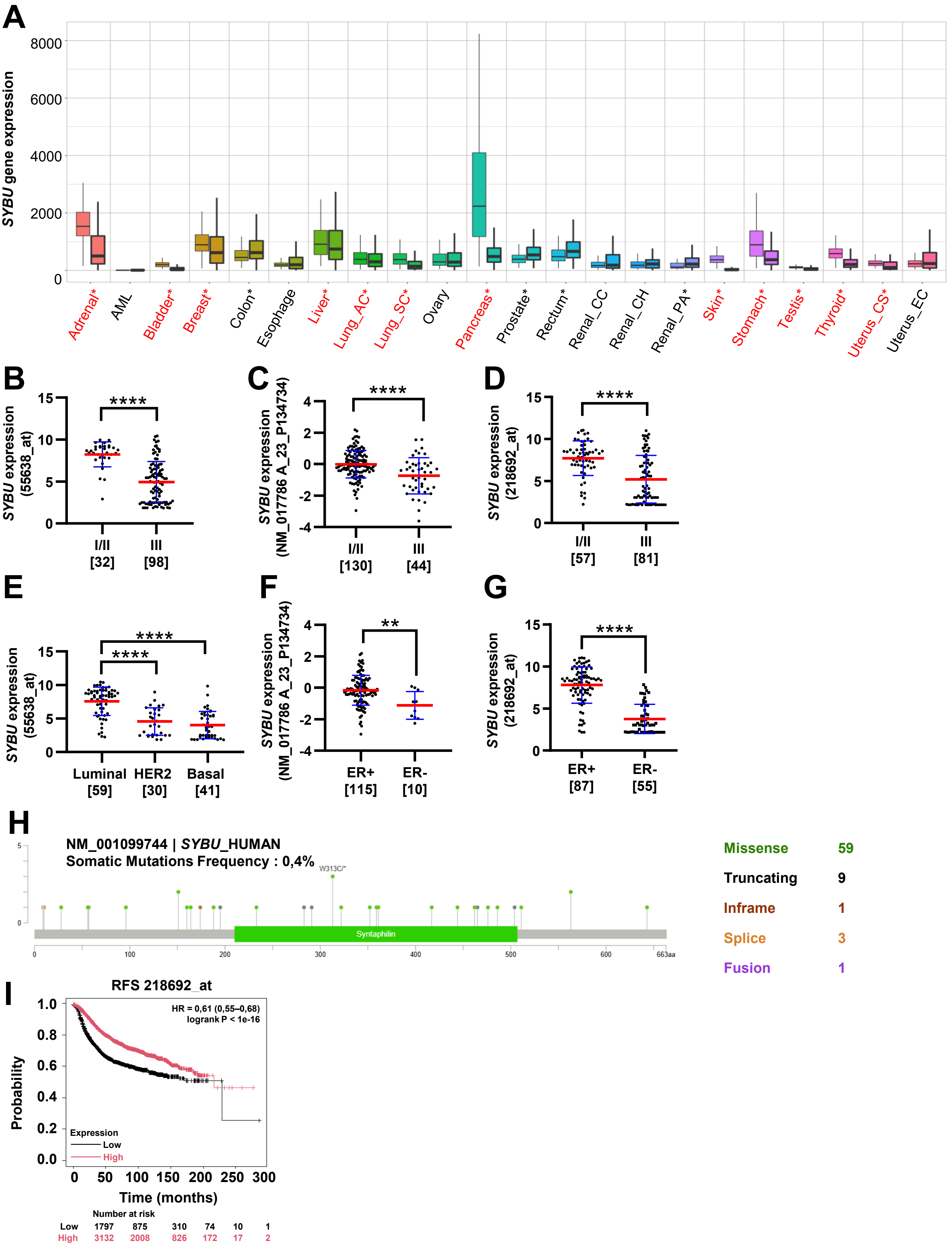


Figure S1 Morin et al.

**Figure S1. *SYBU* expression levels in cancer**

**(A)** RNA-seq analysis of *SYBU* gene expression in normal (left) versus tumor tissues (right) from tnmplot.com. Asterisks indicate significant difference between normal and tumor. *SYBU* downregulation in tumors is indicated in red. AML *Acute Myeloid Leukemia;* AC *Adenocarcinoma*; SC *Squamous Cell Carcinoma*; CC *Clear Cell carcinoma;* CH *Chromophobe carcinoma;* PA *Papillary carcinoma*; CS *Carcinosarcoma*; EC *Endometrioid carcinoma.* **(B-D)** *SYBU* probeset intensity according to histological grade in Curie **(B)**, IGR **(C)** and R04 **(D)** cohorts. **(E)** Same as (B) in breast tumors according to molecular subtypes. **(F,G)** *SYBU* probeset intensity in estrogen receptor (ER) positive and negative tumors from IGR **(F)** and R04 **(G)** cohorts. **(H)** Lollipop plot of somatic mutations in SYBU across all breast cancer datasets from cBioPortal.org (N=17076 samples; N=15404 patients in 32 studies). Each vertical line represents a mutation observed in tumor samples, y axis indicating the number of cases. The colored circles indicate the mutation type. **(I)** Kaplan-Meier survival curves from kmplot.com based on *SYBU* expression (probeset 218692_at) showing relapse-free survival (RFS). The probeset best cutoff was selected.

**Data information.** Number of tumors is indicated under brackets. Statistical analyses: **(B,D,F,G)** Mann-Whitney; **(C)** Unpaired T-test; **(E)** Kruskal-Wallis with Dunn's *post hoc* and **(I)** log-rank tests. p**<0.01, p****<0.0001.

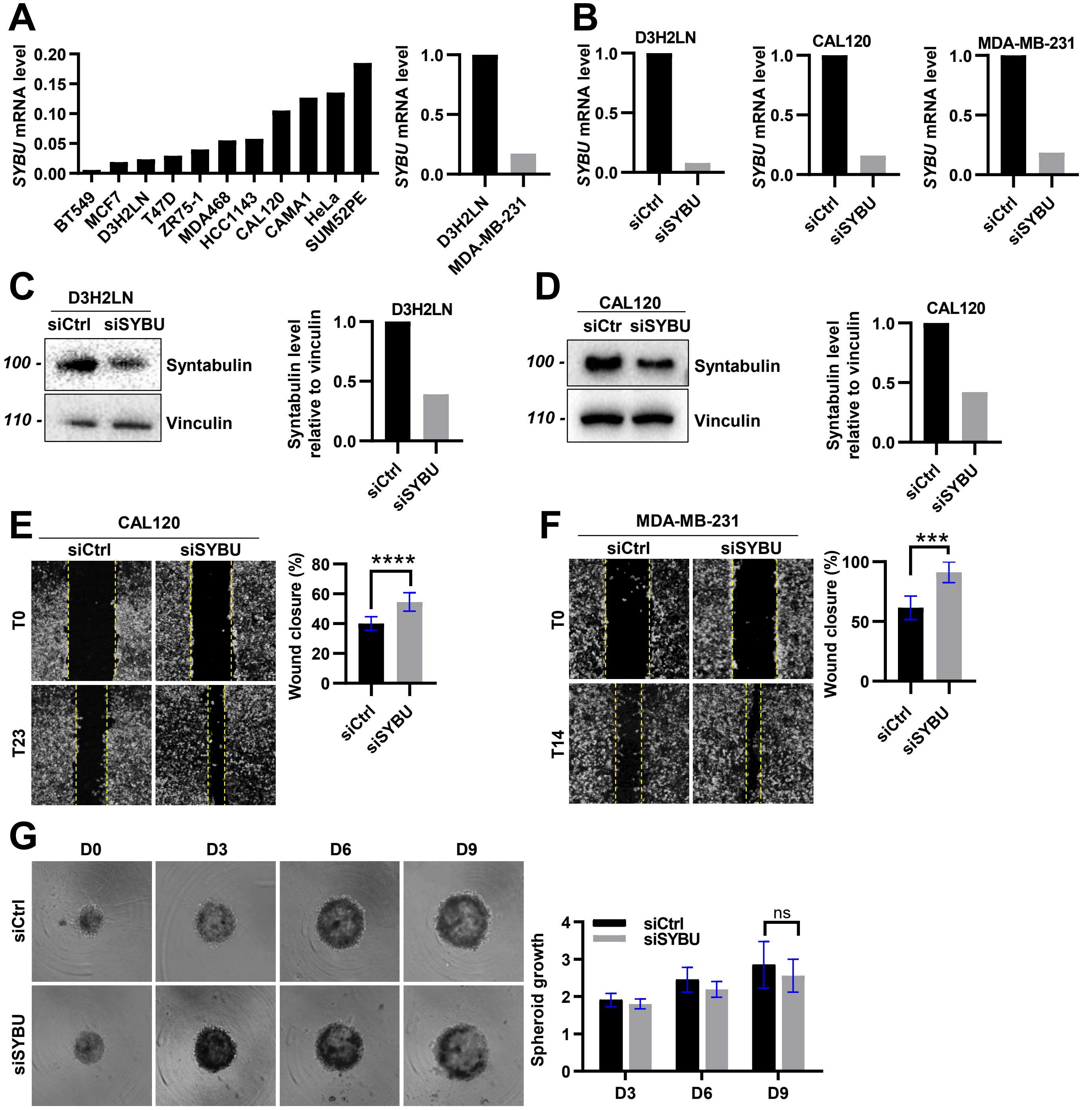
A
SYBU mRNA level
BT549
MCF7
D3H2LN
T47D
ZR75-1
MDA468
HCC1143
CAL120
CAMA1
HeLa
SUM52PE
MDA-MB-231
B
D3H2LN
CAL120
MDA-MB-231
siCtrl
siSYBU
C
D3H2LN
siCtrl siSYBU
100 -
110 -
Syntabulin
Vinculin
Syntabulin level relative to vinculin
D
CAL120
siCtr siSYBU
E
CAL120
T0
T23
Wound closure (%)
****
F
MDA-MB-231
T14
***
G
D0
D3
D6
D9
Spheroid growth
ns

**Figure S2. *SYBU* silencing impairs tumor cell migration and invasion**

**(A)** *SYBU* expression in different breast cancer cell lines relative to *HPRT1* (*left*) and *RPL13* (*right*) genes as assessed by qPCR. **(B)** qPCR analysis of *SYBU* expression as in (A) in siCtrl or siSYBU-transfected D3H2LN, CAL120 and MDA-MB-231 cells. **(C,D)** Immunoblot analysis of syntabulin expression in siCtrl or siSYBU-transfected D3H2LN **(C)** and CAL120 **(D)** cells. *Right:* Quantification of syntabulin relative to vinculin intensity. **(E,F)** Wound healing assay in siCtrl or siSYBU-transfected CAL120 **(E)** or MDA-MB-231 cells **(F)**. *Right*: Quantification of wound closure. **(G)** Spheroid growth in siCtrl or siSYBU-transfected D3H2LN cells monitored over day. *Right*: Quantification of area normalised to T0 area.

**Data information.** Error bars represent mean ± SD in one experiment including n=4 wells/condition **(E,F)** and n=6 spheroids/condition **(G)**. Statistical analyses: **(E,F)** Mann-Whitney; **(G)** Two-way ANOVA with Sidak's multiple comparisons tests. ***p<0.001, p****<0.0001.

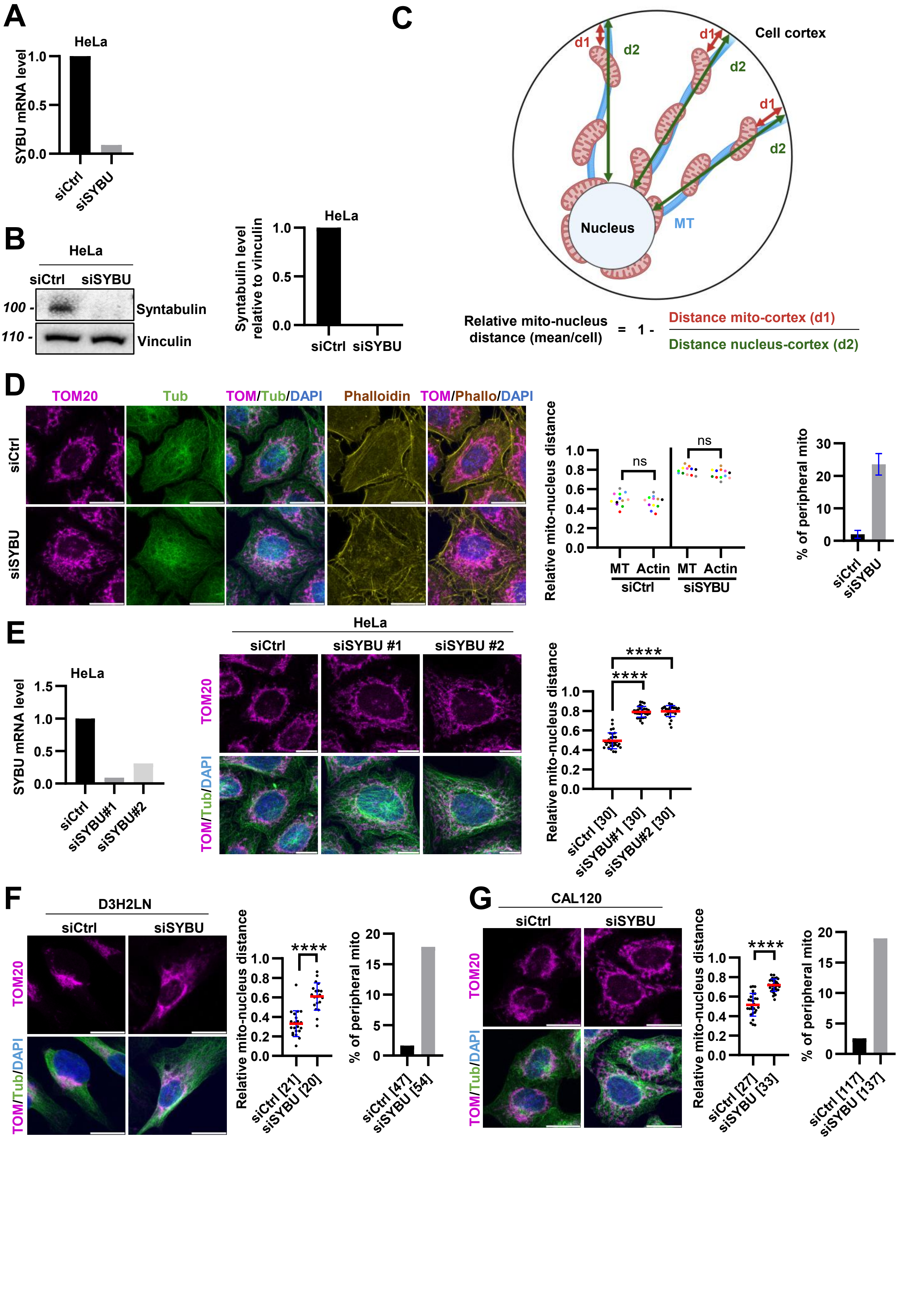


Figure S3 Morin et al.

**Figure S3. *SYBU* depletion enhances anterograde mitochondrial transport along microtubules**

**(A)** qPCR analysis of *SYBU* expression in siCtrl or siSYBU-transfected HeLa cells. **(B)** Immunoblot analysis of syntabulin expression in siCtrl or siSYBU-transfected HeLa cells. *Right:* Quantification of syntabulin relative to vinculin intensity. **(C)** Schematic representation of the approach used to quantify the relative distance between mitochondria and the nucleus (mito-nucleus distance). **(D)** Immunofluorescence of mitochondrial distribution along microtubule (MT) or actin cytoskeleton in siCtrl or siSYBU-transfected HeLa cells. *Middle:* Quantification of relative distance between mitochondria and nucleus, normalised to the distance between nucleus and MT or actin cytoskeleton. *Right:* Percentage of peripheral mitochondria (distance < 2 µm from microtubule end). **(E)** *Left*: qPCR analysis of *SYBU* expression after transfection with siCtrl or two different siSYBU siRNAs (siSYBU#1 and siSYBU#2) in HeLa cells. *Middle*: Immunofluorescence of mitochondrial distribution in siCtrl, siSYBU#1 or siSYBU#2-transfected HeLa cells. *Right:* Quantification of relative distance between mitochondria and nucleus. **(F)** Same as (D) in D3H2LN cells. *Right*: Quantification as (D). **(G)** Same as (D) in CAL120 cells. *Right*: Quantification as in (D).

<u>**Data information**</u>**.** Fluorescent staining shows mitochondria in magenta, tubulin in green and phalloidin in yellow. Shown is one experiment. Error bars represent mean ± SD from one experiment. Statistical analyses: **(E)** Kruskal-Wallis with Dunn's *post hoc*; **(F)** Mann-Whitney and **(G)** Unpaired T-tests. p****<0.0001. Number of cells is indicated under brackets. Scale bar = 10 µm.

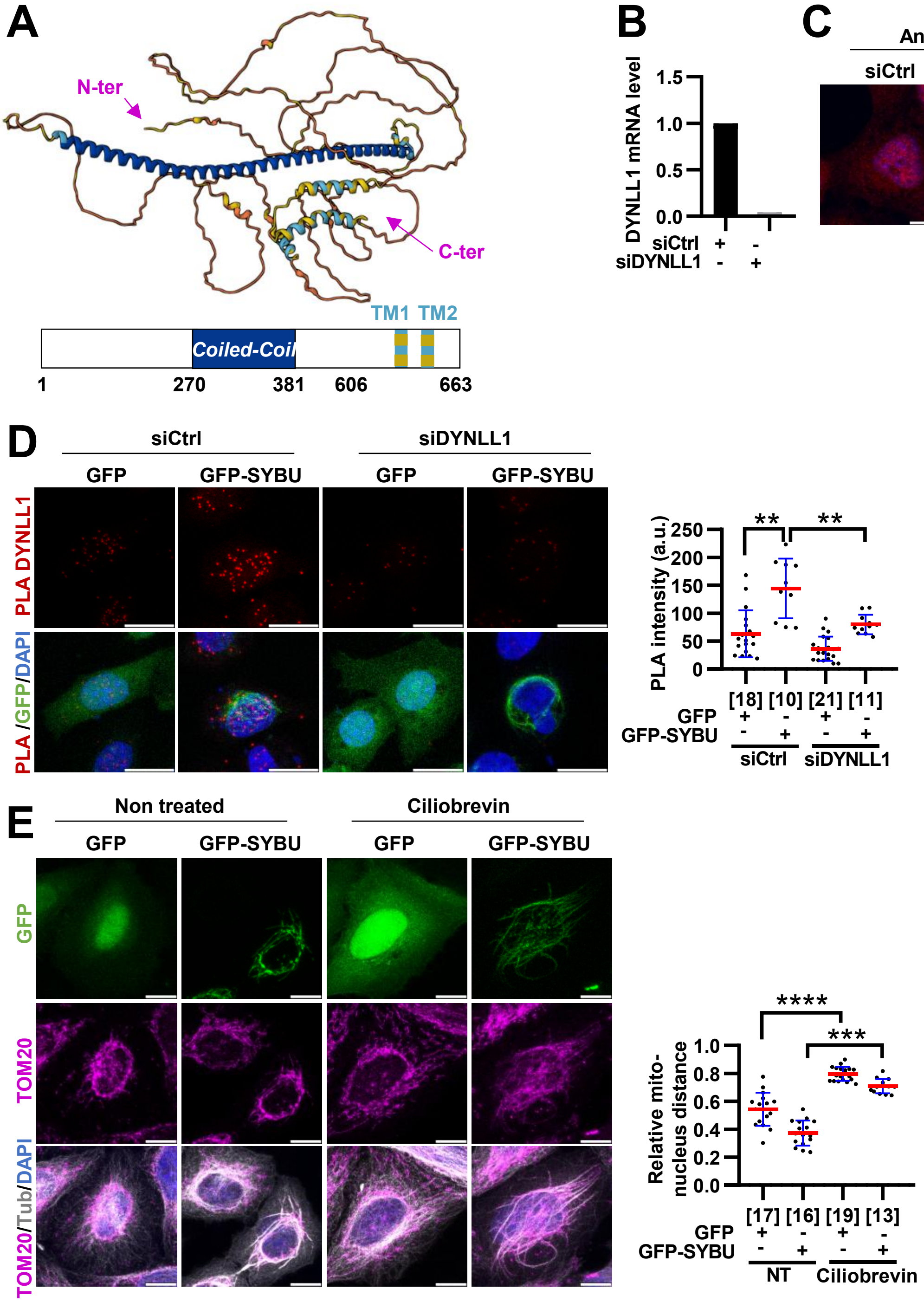


Figure S4 Morin et al.

**Figure S4. Syntabulin interacts with dynein light chain 1 DYNLL1**

**(A)** Predicted 3-dimensional structure of syntabulin obtained using AlphaFold, showing long disordered segments at the C-terminus, an extended *coiled-coil* dimerization domain, and two C-terminal transmembrane α-helices. **(B)** qPCR analysis of DYNLL1 expression in siCtrl or siDYNLL1-transfected HeLa cells. **(C)** Immunofluorescence of anti-DYNLL1 in siCtrl or siDYNLL1-transfected HeLa cells. *Right*: Quantification of signal intensity. **(D)** Proximity ligation assay (PLA) using anti-DYNLL1 and anti-GFP antibodies in siCtrl or siDYNLL1 HeLa cells expressing GFP or GFP-SYBU. Red puncta indicate proximity between GFP-SYBU and DYNLL1. *Right:* Quantification of rolling circles PLA signal intensity. **(E)** Immunofluorescence of mitochondrial distribution in GFP or GFP-SYBU-transfected HeLa cells treated or not with 25 μM of Ciliobrevin D. *Right*: Quantification of relative distance between mitochondria and nucleus.

**Data information.** Fluorescent staining shows **(C)** DYNLL1 in red; **(D)** PLA in red; **(E)** TOM20 in magenta and tubulin in grey. Error bars represent mean ± SD. Shown is one experiment. Statistical analyses: **(C)** Unpaired T-test; **(D,E)** Kruskal-Wallis with Dunn's *post hoc* test. $p^{**}<0.01$, $p^{****}<0.0001$. Number of cells is indicated under brackets. Scale bar = 10 μm.

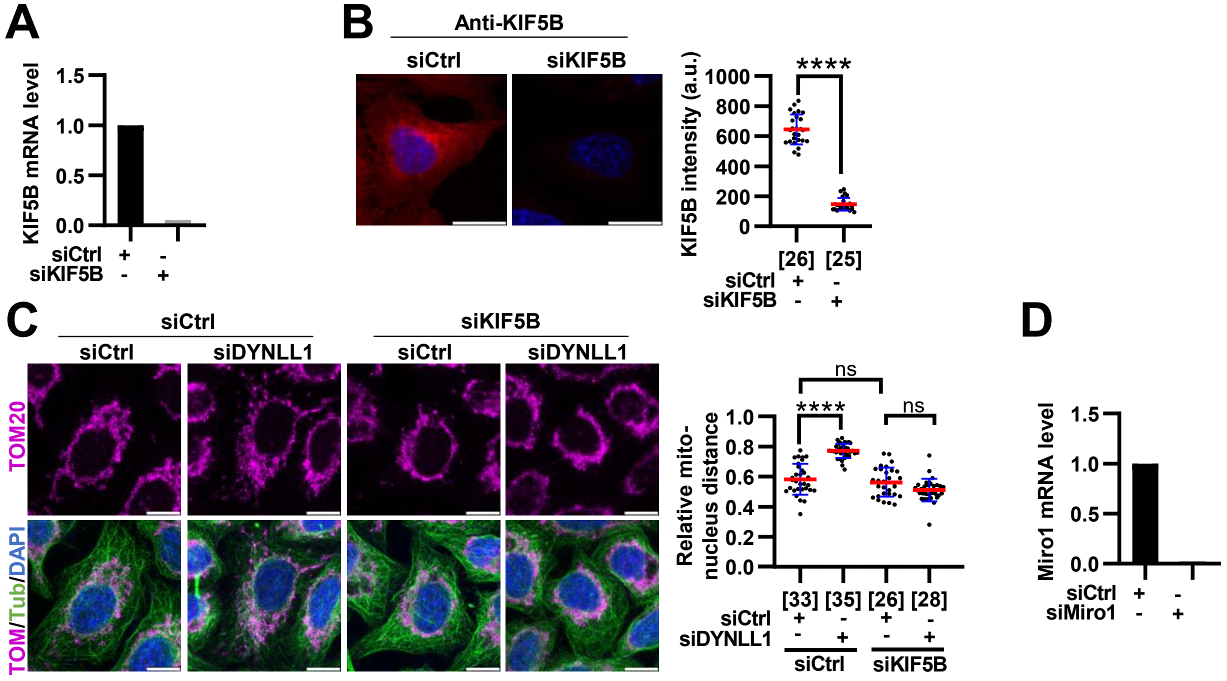


**Figure S5 : Syntabulin opposes Miro/KIF5B-mediated anterograde mitochondrial transport**

**(A)** qPCR analysis of KIF5B expression in siCtrl or siKIF5B-transfected HeLa cells. **(B)** Immunofluorescence of anti-KIF5B in siCtrl or siKIF5B-transfected HeLa cells. *Right*: Quantification of signal intensity. **(C)** Immunofluorescence of mitochondria distribution in siCtrl, siDYNLL1 or siKIF5B-transfected HeLa cells. *Right:* Quantification of relative distance between mitochondria and nucleus. **(D)** Same as (A) with Miro1 depletion.

**Data information.** Fluorescent staining shows : **(B)** KIF5B in red; **(C)** TOM20 in magenta and microtubules in green. Error bars represent mean ± SD from two independent experiments **(C)**. Shown is one experiment **(B)**. Statistical analyses: **(B)** Mann-Whitney and **(C)** Kruskal-Wallis with Dunn's *post hoc* tests. p****<0.0001. Number of cells is indicated under brackets. Scale bar = 10 µm.

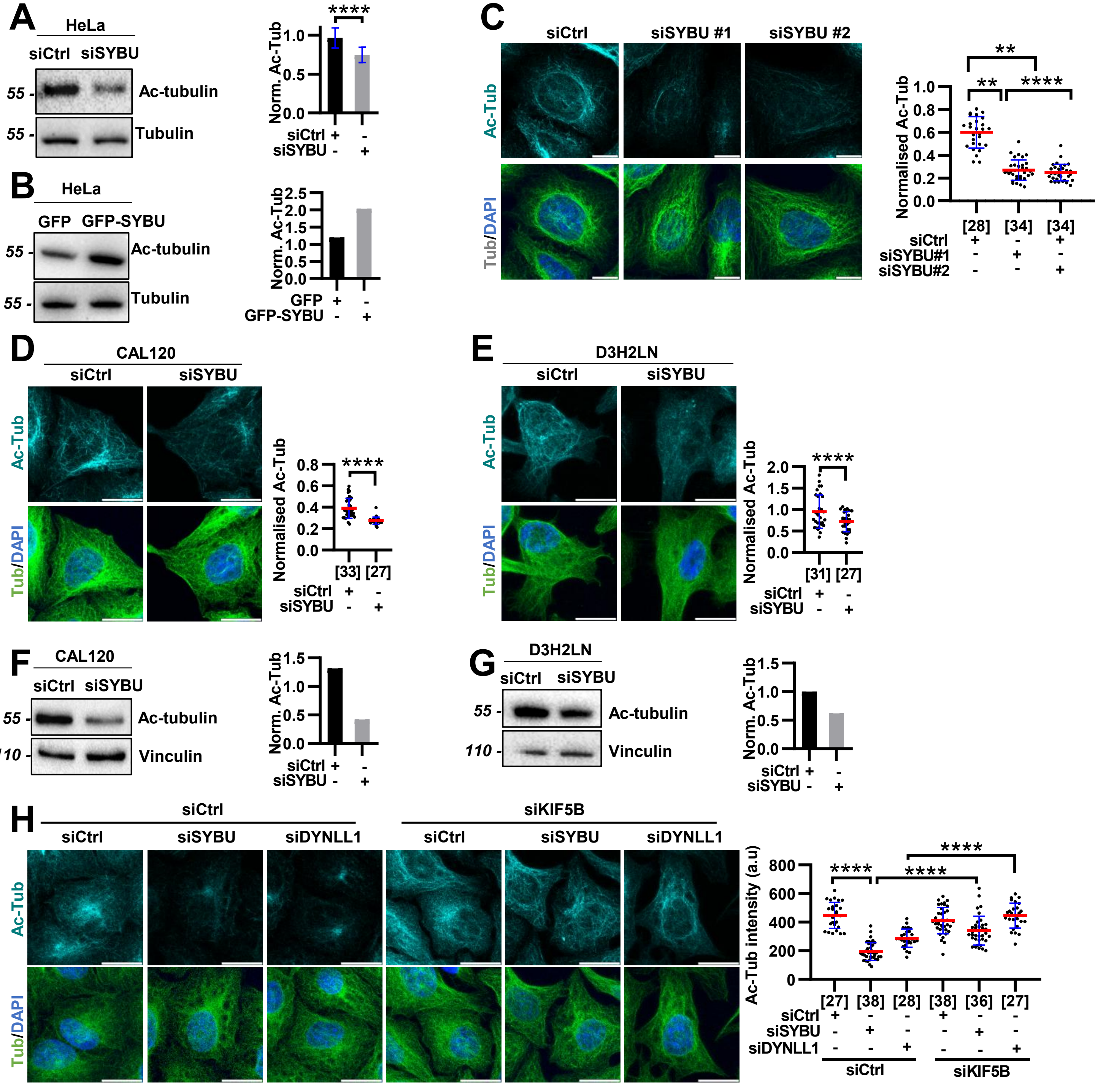


Figure S6 Morin et al.

**Figure S6. *SYBU* depletion decreases microtubule acetylation and causes microtubule damage**

**(A)** Immunoblotting of siCtrl or siSYBU-transfected HeLa cells probed with anti-acetyl-tubulin (Ac-Tub) and anti-tubulin (Tub) antibodies. *Right:* Quantification of Ac-Tubulin relative to Tub intensity. **(B)** Same as (A) in GFP-SYBU or GFP-transfected HeLa cells. *Right:* Quantification as (A). **(C)** Immunofluorescence of microtubule acetylation in siCtrl, siSYBU#1 or siSYBU#2-transfected HeLa cells. *Right:* Quantification of Ac-Tub relative to Tub intensity. **(D,E)** Immunofluorescence of microtubule acetylation in siCtrl or siSYBU-transfected CAL120 **(D)** or D3H2LN cells **(E)**. *Right:* Quantification as (C). **(F,G)** Same as (A) in CAL120 **(F)** or D3H2LN cells **(G)**. *Right:* Quantification as (A). **(H)** Immunofluorescence of microtubule acetylation in siCtrl, siSYBU, siKIF5B or siDYNLL1-transfected HeLa cells. *Right:* Quantification of Ac-Tub intensity.

**Data information.** Fluorescent staining shows Ac-Tub in cyan and Tub in green. Error bars represent mean ± SD from **(A)** 11 independent experiment. Shown is one experiment **(B-H)**. Statistical analyses: **(A,D,E)** Mann-Whitney; **(C,H)** Kruskal-Wallis with Dunn's *post hoc* tests. $p^* < 0.05$, $p^{****} < 0.0001$. Number of cells is indicated under brackets. Scale bar = 10 µm.

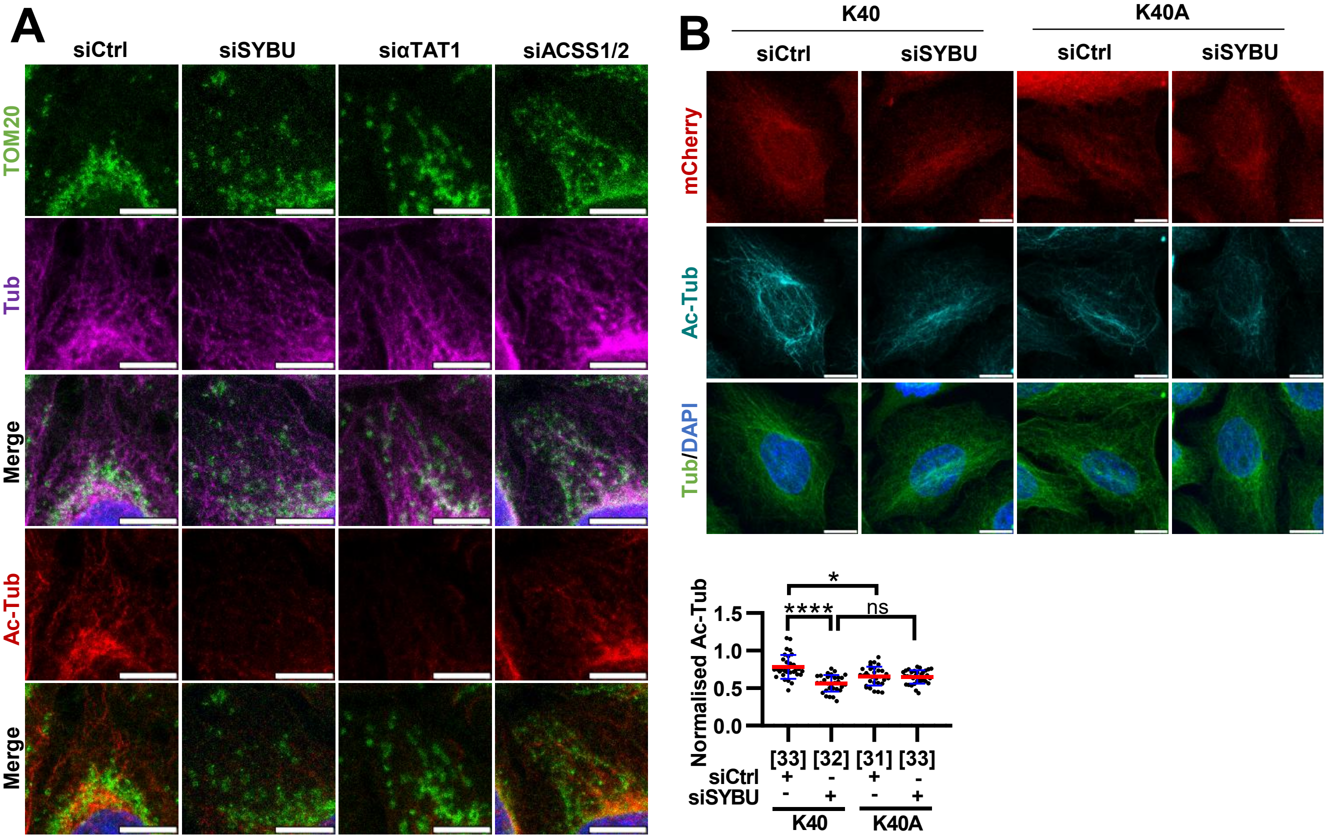


**Figure S7. Microtubule deacetylation promotes anterograde mitochondrial transport**

**(A)** Cropped images of Fig. 6A showing mitochondria distribution in siCtrl, siSYBU, siαTAT1 or siACSS1/2-transfected HeLa cells. **(B)** Immunofluorescence of Ac-Tub in siCtrl or siSYBU-transfected HeLa cells expressing mCherry-K40K (wild type α-tubuline K40 acetylation) or mCherry-K40A (non-acetylable). *Bottom:* Quantification of Ac-Tub relative to Tub intensity.

**Data information**. Fluorescent staining shows **(A)** mitochondria in green, Tub in magenta and Ac-Tub in red; **(B)** mCherry in red, Ac-Tub in cyan, Tub in green. Error bars represent mean ± SD. Shown is one experiment. Statistical analyses: **(B)** One-way ANOVA with Tukey's *post hoc* test was performed. p****<0.0001. Number of cells is indicated under brackets. Scale bar = 5 µm **(A)** ; 10 µm **(B)**.

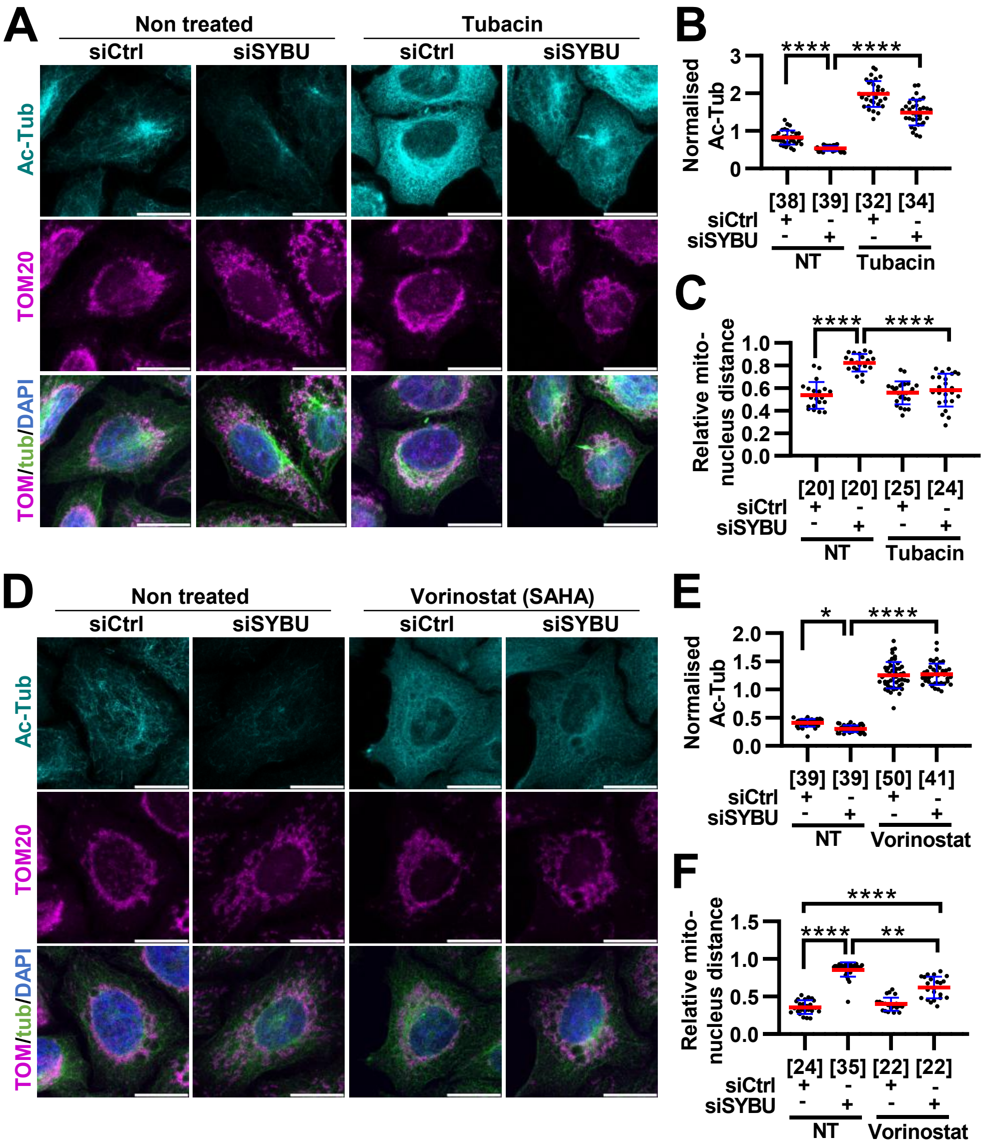


**Figure S8. HDAC6 inhibition rescues *SYBU* silencing phenotype**

**(A)** Immunofluorescence of mitochondria distribution and microtubule acetylation in siCtrl or siSYBU-transfected HeLa cells, treated or not with Tubacin (1 µM, 4 hrs, 37°). **(B)** Quantification from (A) of Ac-Tub relative to Tub intensity. **(C)** Quantification from (A) of relative distance between mitochondria and nucleus. **(D)** Same as (A) with Vorinostat treatment or not (1 µM, 4 hrs, 37°). **(E)** Quantification from (D) as (B). **(F)** Quantification from (D) as (C).

**Data information**. Fluorescent staining shows Ac-Tub in cyan, mitochondria in magenta and Tub in green. Error bars represent mean ± SD from two independent experiment **(A-F)**. Statistical analyses: **(B,C)** One-way ANOVA with Tukey's *post hoc;* **(E,F)** Kruskal-Wallis with Dunn's *post hoc* tests. p*<0.05, p**<0.01, p****<0.0001. Number of cells is indicated under brackets. Scale bar = 10 µm.

### Supplementary Videos

**Supplementary video 1**: Five-minute time-lapse videomicroscopy showing mitochondrial transport (MitoTracker in magenta) in D3H2LN-GFP cells (green) transfected with control siRNA, recorded 30 minutes post-seeding.

**Supplementary video 2**: Five-minute time-lapse videomicroscopy showing mitochondrial transport (MitoTracker in magenta) in D3H2LN-GFP cells (green) transfected with siRNA targeting syntabulin (siSYBU), recorded 30 minutes post-seeding.
